# A septin GTPase scaffold of dynein-dynactin motors triggers retrograde lysosome transport

**DOI:** 10.1101/2020.06.01.128488

**Authors:** Ilona A. Kesisova, Benjamin P. Robinson, Elias T. Spiliotis

## Abstract

The metabolic and signaling functions of lysosomes depend on their intracellular positioning and trafficking, but the underlying mechanisms are little understood. Here, we have discovered a novel septin GTPase-based mechanism for retrograde lysosome transport. We found that septin 9 (SEPT9) associates with lysosomes, promoting the perinuclear localization of lysosomes in a Rab7-independent manner. SEPT9 targeting to mitochondria and peroxisomes is sufficient to recruit dynein and cause perinuclear clustering. We show that SEPT9 interacts with both dynein and dynactin through its GTPase domain and N-terminal extension, respectively. Strikingly, SEPT9 associates preferentially with the dynein intermediate chain (DIC) in its GDP-bound state, which favors dimerization and assembly into septin multimers. In response to oxidative cell stress induced by arsenite, SEPT9 localization to lysosomes is enhanced, promoting the perinuclear clustering of lysosomes. We posit that septins function as GDP-activated scaffolds for the cooperative assembly of dynein-dynactin, providing an alternative mechanism of retrograde lysosome transport at steady state and during cellular adaptation to stress.

**Summary:** The intracellular position of lysosomes is critical for cell metabolism and signaling. Kesisova et al discovered a membrane-associated septin GTPase scaffold of dynein-dynactin that promotes retrograde traffic and perinuclear lysosome clustering at steady state and in response to oxidative stress.

## Introduction

Lysosomes are major degradative organelles with critical functions in a diversity of cellular processes including cell metabolism, signaling, gene regulation and immunity (Blott and Griffiths, 2002; Lawrence and Zoncu, 2019; Settembre et al., 2013). Lysosomes contain membrane transporters of amino acids, nucleotides, lipids and ions, which sense intracellular conditions and crosstalk with signaling complexes that regulate autophagy and gene transcription (Li et al., 2019; Lim and Zoncu, 2016; Schwake et al., 2013). Lysosomes are dynamic organelles, whose intracellular position and movement are critical for their signaling functions, maturation, turnover and interaction with other membrane organelles (Bonifacino and Neefjes, 2017; Luzio et al., 2007; Saftig and Klumperman, 2009; Savini et al., 2019). In response to nutritional and oxidative stress, lysosomes mobilize to perinuclear areas of the cytoplasm, where they fuse with autophagosomes (Lim and Zoncu, 2016; Yim and Mizushima, 2020). Similarly, lysosomes traffic retrogradely to bacteria undergoing autophagy (Hu et al., 2020) and anterogradely to plasma membrane sites of repair (Andrews and Corrotte, 2018), and exocytose in migrating and immune cells (Castro-Castro et al., 2016; Lettau et al., 2007; Wilson et al., 2018).

How lysosomes mobilize in response to various intracellular conditions and cues is not well understood. Lysosome positioning and movement involve selective association with microtubule motors and subsets of microtubules with distinct post-translational modifications (Bonifacino and Neefjes, 2017; Guardia et al., 2016; Mohan et al., 2019). Anterograde movement of lysosomes to the cellular periphery is mediated by motors of the kinesin-1, -2 and -3 families (Farias et al., 2017; Guardia et al., 2016; Matsushita et al., 2004; Pankiv et al., 2010; Rosa-Ferreira and Munro, 2011). Retrograde movement of lysosomes to the perinuclear cytoplasm is driven by the microtubule motor dynein. Association of dynein with lysosomes occurs through mechanisms which are linked to the metabolic sensing functions of lysosomes. Components of the dynein-dynactin complex interact directly with the calcium ion sensor ALG2 and the cholesterol-sensing Rab7-RILP-ORP1L complex (Li et al., 2016; Rocha et al., 2009). Additionally, dynein associates with the scaffold protein JIP4, which is recruited by the lysosomal transmembrane protein TMEM55B, whose levels are upregulated in response to lysosomal stress (Willett et al., 2017). Despite this multimodal recruitment of dynein-dynactin, which indicates that lysosomes adopt a diversity of strategies for retrograde transport, it is little understood how dynein motility is activated on lysosomal membranes.

The discovery of adaptor proteins that promote the interaction of dynein with dynactin has revolutionized our understanding of dynein motility (Cross and Dodding, 2019; McKenney et al., 2014; Olenick and Holzbaur, 2019; Reck-Peterson et al., 2018). Dynein is a hexameric motor consisting of a dynein heavy chain (DHC), an intermediate chain (DIC), a light intermediate chain (DLIC) and three light chains (Schmidt and Carter, 2016; Sweeney and Holzbaur, 2018). Dynactin is a large multi-subunit complex that is made of a central actin-like filament, which is capped by proteins on its barbed (CAPZ) and pointed ends (ARP11, p62, p27, p25), and a shoulder sub-complex containing p150^GLUED^, p50 dynamitin and p24 (Schroer, 2004). Dynein dimerizes into an autoinhibitory conformation, which is weakly processive and requires assembly with dynactin and activating adaptor proteins in order to move efficiently on microtubules (Chowdhury et al., 2015; McKenney et al., 2014; Schroer and Sheetz, 1991; Urnavicius et al., 2018; Zhang et al., 2017). Dynein adaptors contain coiled-coil domains that run along the central dynactin filament and promote dynein-dynactin binding by making contacts with both dynactin and the dynein heavy and light intermediate chains (Chowdhury et al., 2015; Lee et al., 2018; Olenick and Holzbaur, 2019; Schlager et al., 2014; Schroeder and Vale, 2016; Urnavicius et al., 2018). These coiled coil domains, however, are not present in all dynein adaptors including the lysosomal RILP and JIP4 (Reck-Peterson et al., 2018).

Septins are a family of GTP-binding proteins, which multimerize into higher order oligomers and polymers that associate with cell membranes and the cytoskeleton (Mostowy and Cossart, 2012; Spiliotis, 2018). Membrane-bound septins function as scaffolds and diffusion barriers, which control protein localization in a spatiotemporal specific manner (Bridges and Gladfelter, 2015; Caudron and Barral, 2009). In the endocytic pathway, septins associate preferentially with endolysosomes that are enriched with phosphatidylinositol 3,5-bisphosphate and Rab7 (Dolat and Spiliotis, 2016). Moreover, septins are critical for lysosome merging with macropinosomes and *Shigella* bacteria undergoing autophagy (Dolat and Spiliotis, 2016; Krokowski et al., 2018). In a proteomic study of the dynein interactome, septins were identified as potential binding partners of DIC, DLIC and the dynein adaptor BiCD2 (Redwine et al., 2017). Here, we report that membrane-associated septins provide a novel GDP-activated mechanism for the retrograde dynein-driven transport of lysosomes.

## Results

### Septins associate with lysosomes and promote retrograde trafficking in a Rab7-independent manner

Septins have been reported to associate with membranes of the late endocytic pathway, localizing to mature macropinosomes and impacting the formation of multivesicular bodies (Dolat and Spiliotis, 2016; Traikov et al., 2014). However, it is unknown whether septins are functionally present on lysosomes. Using high- and super-resolution microscopy as well as subcellular fractionation, we probed for septin localization to lysosomes. We stained COS-7 cells with antibodies against septin 9 (SEPT9), a ubiquitously expressed septin and core subunit of heteromeric septin complexes (Kim et al., 2011; Sellin et al., 2011). As previously observed (Connolly et al., 2011; Verdier-Pinard et al., 2017), SEPT9 formed stress fiber-like filaments on the ventral side of the nucleus and localized to plasma membrane domains of curvature. Additionally, SEPT9 puncta were visible in perinuclear and peripheral regions of the cytoplasm. Confocal microscopy showed that a fraction of lysosomes and early endosomes, which were immunolabeled with antibodies against LAMTOR4 and EEA1, respectively, overlapped with SEPT9 filaments and puncta (Fig. 1 A-B). Quantitatively, SEPT9 puncta were present on lysosomes more abundantly than early endosomes (79.7% vs. 40.8%) indicating a preferential association with lysosomes (Fig. 1 C). SEPT9 puncta, however, covered 23% ± 0.4% of the total surface of lysosomes compared to 12% ± 0.4% of early endosomes per cell (*n* = 6), suggesting that SEPT9 localizes to subdomains of the lysosomal membrane. Super-resolution structured illumination microscopy showed that SEPT9 puncta localize to microdomains of the limiting membrane of LAMP2-positive compartments (Fig. 1 D). These SEPT9 microdomains resembled the clusters of LAMP2A and dynein motors, which have been previously observed on the membranes of late phagosomes and lysosomes (Kaushik et al., 2006; Rai et al., 2016). In living cells, SEPT9-mCherry (SEPT9 isoform 1), which was expressed under the weak PGK promoter, also localized to microdomains of LAMP1-mGFP compartments (Fig. 1 E, arrows) and co-trafficked with LAMP1-mGFP compartments that moved retrogradely toward the nucleus (Fig. 1 F, Video 1). In contrast, colocalization and co-traffic of SEPT9-mCherry with GFP-EEA1 was rare (Fig. S1 A-B) Consistent with septin localization to lysosomes, septins (SEPT2, SEPT6, SEPT7, SEPT9) cofractionated with LAMP1- and Rab7-positive membranes at the top of an iodixanol (Optiprep™) density gradient of human embryonic kidney cell (HEK293) extracts (Fig. 1 G). In the top endolysosomal Rab7- and Rab5-containing fractions, SEPT9 exhibited a distinct co-enrichment with LAMP1 (fraction #1) compared to SEPT2, SEPT6 and SEPT7, which were more widely distributed (Fig. 1 G).

**Figure 1.**
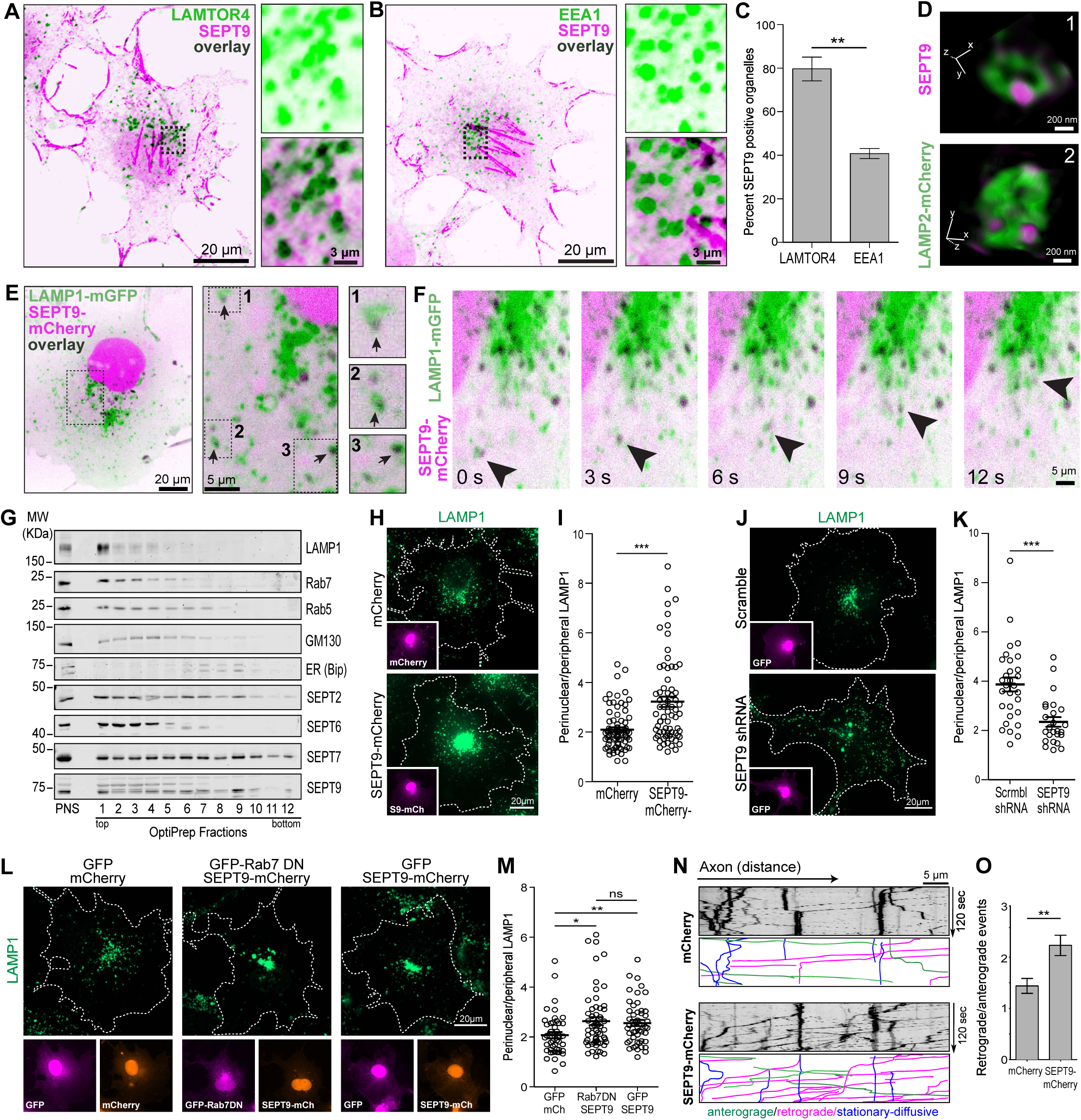
Septins associate with lysosomes and promote positioning in the perinuclear cytoplasm. (A-B) Confocal images of COS-7 cells stained for SEPT9 (inverted magenta) and LAMTOR4 (inverted green; A) or EEA1 (inverted green; B). Insets show lysosomes from perinuclear regions in higher magnification. (C) Bar graph shows percentage (mean ± SEM) of LAMTOR4- and EEA1-organelles with SEPT9 puncta per cell (*n* = 6). (D) Super-resolution SIM images show SEPT9 puncta on the delimiting membrane of LAMP2-mCherry labeled compartments in BSC-1 cells. (E-F) Spinning disk confocal microscopy image (E) and still frames from time-lapse imaging (F) of COS-7 cells transfected with SEPT9_i1-mCherry and LAMP1-mGFP. Insets show in higher magnification LAMP1-mGFP compartments with SEPT9_i1-mCherry puncta (arrows), and arrowheads point to retrograde co-traffic toward the nucleus. (G) Post-nuclear supernatants (PNS) of HEK-293 cell extracts were loaded on an iodixanol OPTIPREP density gradient (17%-20%-23%-27%-30%) and fractions were blotted for LAMP-1, Rab7, Rab7, GM130, BIP, SEPT9, SEPT2, SEPT6 and SEPT7. (H) COS-7 cells were transfected with mCherry or SEPT9-mCherry (insets) and stained with anti-LAMP1. (I) Plot shows the ratio (mean ± SEM) of perinuclear to peripheral LAMP1 fluorescence intensity per cell (*n* = 70-79). (J) COS-7 were treated with control or SEPT9-targeting shRNAs and stained with anti-LAMP1. (K) Plot shows the ratio (mean ± SEM) of perinuclear to peripheral LAMP1 fluorescence intensity per cell (*n* = 25-30). (L) Images show lysosome (LAMP-1) distribution in COS-7 cells transfected with mCherry/GFP, SEPT9-mCherry/GFP-Rab7-DN and SEPT9-mCherry/GFP. (M) Quantification shows the ratio (mean ± SEM) of perinuclear to peripheral LAMP1 fluorescence intensity per cell (*n* = 45-51). (N) Embryonic (E18) hippocampal neurons (DIV4) were transfected with LAMP-1-mGFP and mCherry or rat SEPT9-mCherry, and axons imaged live by TIRF microscopy. Kymographs show stationary or diffusive (blue), retrogradely (magenta) or anterogradely (green) moving particles with LAMP1-mGFP. (O) Bar graph shows the ratio (mean ± SEM) of retrogradely to anterogradely moving LAMP-1-mGFP particles per 2 minutes per axon (*n* = 18-19). Ns, non-significant (p > 0.05); *, p < 0.05; **, p < 0.01; *** p < 0.001.

To test whether septins impact the intracellular position of lysosomes, we over-expressed SEPT9 in COS-7 cells. Strikingly, SEPT9-mCherry increased the perinuclear clustering of lysosomes, increasing the ratio of perinuclear to peripheral lysosomes (Fig. 1 H-I); high levels of nuclear and cytoplasmic SEPT9-mCherry concealed its lysosomal localization, which was observed in cells expressing SEPT9-mCherry under a weak promoter (Fig. 1 E-F). In contrast to the perinuclear clustering of lysosomes, the intracellular distribution of early endosomes did not change (Fig. S1 C-D), and over-expression of SEPT2 and SEPT6 did not impact lysosomal distribution (Fig. S1 E-F). Depletion of SEPT9 diminished the perinuclear population of lysosomes, exerting the opposite effect of SEPT9 over-expression (Fig. 1 J-K and S1 G-H).

Given that retrograde dynein-driven transport of lysosomes is hitherto controlled by the small GTPase Rab7, we asked if Rab7 is required for the perinuclear clustering of lysosomes by SEPT9. Using a Rab7 dominant negative (Rab7-DN) mutant, we quantified the intracellular distribution of lysosomes in COS-7 cells, which over-expressed SEPT9 in the presence or absence of Rab7-DN. Rab7-DN did not alter the effects of SEPT9 over-expression, indicating that SEPT9 induces retrograde lysosome transport independently of Rab7 (Fig. 1 L-M).

Next, we sought to examine whether SEPT9 has a conserved role in lysosome positioning across different cell types. We tested if SEPT9 impacts lysosome traffic in primary rat embryonic hippocampal neurons. In early stages of neuronal morphogenesis, SEPT9 is present in the developing axon before it becomes enriched on dendritic microtubules (Karasmanis et al., 2018). In the axons of these cells, which consist of unidirectional plus end-out oriented microtubules, we found that SEPT9 was enriched more than two-fold in LAMP1-positive endolysosomes compared to Rab5-containing early endosomes (Fig. S1 I-J). SEPT9 over-expression enhanced the retrograde trafficking events of axonal lysosomes, increasing the ratio of retrograde-to-anterograde events from 1.44 ± 0.15 to 2.23 ± 0.20 (Fig. 1 N-O, Videos 2 and 3). Taken together with our observations in COS-7 cells, these data show that septins have a functional role in lysosome positioning and trafficking.

### Membrane-associated SEPT9 induces retrograde trafficking by recruiting the dynein-dynactin complex

Because the perinuclear localization of lysosomes depends on SEPT9 expression, we hypothesized that SEPT9 induces retrograde lysosome transport by recruiting dynein-dynactin motors. To directly test this possibility, we generated SEPT9 chimeras that are coupled to the membrane of peroxisomes and mitochondria. Using the rapamycin-inducible FRB-FKBP dimerization system, we targeted GFP-SEPT9-FRB to peroxisomes containing PEX-RFP-FKBP. After 45 minutes of rapalog treatment, GFP-SEPT9-FRB colocalized with PEX-RFP-FKBP and peroxisomes became clustered into a focal juxtanuclear region (Fig. 2 A). Quantification of the percentage of cells with a clustered peroxisome phenotype showed an increase from 35% to 81% after treatment with rapalog (Fig. 2 A). We observed similar perinuclear clustering of mitochondria upon expression of SEPT9-MitoTagRFP, a SEPT9 chimera that contains the mitochondrial-targeting signal of the *Listeria monocytogenes* protein ActA (Fig. 2 B). Perinuclear mitochondria were stained positively for the MitoSpy™ NIR DiIC1 dye, which labels mitochondria with a healthy membrane potential, indicating that the perinuclear aggregation of mitochondria was not due to mitophagy (Fig. S2 A-B).

**Figure 2.**
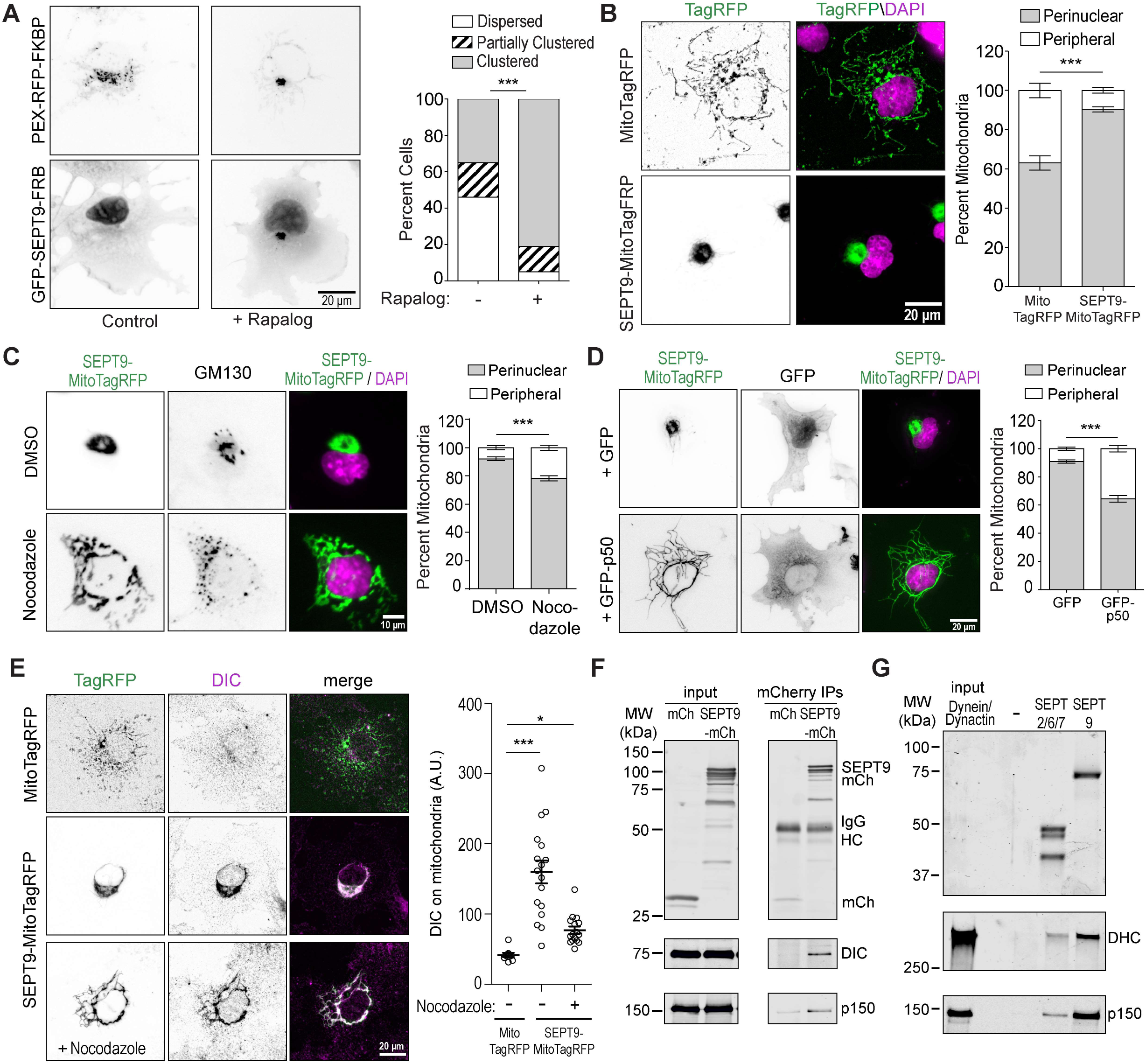
Membrane-associated SEPT9 recruits dynein and induces retrograde dynein-driven transport. (A) Images show COS-7 cells that were transfected with GFP-SEPT9-FRB and PEX-RFP-FKBP after 45 minutes of treatment with paralog or control carrier. Bar graphs shows percentage of cells (*n* = 37-42) with dispersed, partially clustered and clustered peroxisomes. (B) COS-7 cells were transfected with MitoTagRFP or SEPT9-MitoTagRFP and stained with DAPI. Bar graph shows the amount of perinuclear and peripheral mitochondria as percentage (mean ± SEM) of total mitochondria per cell (*n* = 20-44). (C) COS-7 cells were transfected with Mito-TagRFP or SEPT9-MitoTagRFP, treated with nocodazole (33 μM) or DMSO for 3 h, and stained with anti-GM130 and DAPI. Quantification shows the amount of perinuclear and peripheral mitochondria as percentage (mean ± SEM) of total mitochondria per cell (*n* = 53-54). (D) Images show DAPI-stained COS-7 cells that were transfected with SEPT9-MitoTagRFP- and GFP (control) or GFP-p50 (dynamitin). Bar graph shows quantification of perinuclear and peripheral mitochondria as percentage (mean ± SEM) of total per cell (*n* = 55-70). (E) COS-7 cells were transfected with MitoTagRFP or SEPT9-MitoTagRFP and stained with anti-DIC and DAPI after treatment with DMSO or nocodazole (33 μM, 3 h). Plot shows DIC fluorescence intensity (mean ± SEM) on MitoTagRFP-labelled mitochondria per cell (*n* = 9-16). (F) Lysates of HEK-293 cells expressing mCherry or SEPT9-mCherry were incubated with anti-mCherry, and immunoprecipitates were blotted with antibodies against mCherry (top), DIC and p150^GLUED^. (G) Coomassie-stained gel (top) shows recombinant SEPT9 and SEPT2/6/7, which were used in pull-down assays with dynein-dynactin purified from HEK-293 cells. Western blots (bottom) were performed with antibodies against DHC and p150^GLUED^. *, p < 0.05; **, p < 0.01; *** p < 0.001.

Next, we examined whether perinuclear clustering of mitochondria was due to microtubule-dependent dynein-driven transport. Depolymerization of microtubules with nocodazole and expression of the GFP-p50-dynamitin, which inhibits dynein-driven transport in a dominant negative manner (Burkhardt et al., 1997), diminished the perinuclear aggregation of mitochondria resulting in a more dispersed localization throughout the cytoplasm (Fig. 2 C-D). Notably, SEPT9-MitoTagRFP increased the mitochondrial levels of DIC, which remained above control levels upon microtubule depolymerization, indicating that SEPT9 recruits dynein to mitochondria independently of microtubule attachment (Fig. 2 E). Thus, mitochondria-targeted SEPT9 induce retrograde microtubule-dependent transport by promoting the recruitment of dynein.

To further test whether SEPT9 interacts with dynein, we performed co-immunoprecipitations and protein-binding assays. In HEK-293 cell lysates, SEPT9 co-immunoprecipitated with the dynein and dynactin subunits DIC and p150^GLUED^ (Fig. 2 F). Using native dynein and dynactin, which were isolated from HEK-293 cells, we tested whether recombinant SEPT9 interacted directly with dynein. SEPT9 pulled down DHC (dynein) and p150^GLUED^ (dynactin), while binding to SEPT2/6/7 was negligible (Fig. 2 G). Despite lack of SEPT2/6/7-dynein binding, targeting of SEPT2, SEPT6 or SEPT7 to mitochondria enhanced perinuclear clustering (Fig. S2 C-D), indicating that SEPT9 may function in complex with SEPT2/6/7 as mitochondria-targeted SEPT9 also recruited SEPT2, SEPT6 and SEPT7 to mitochondria (Fig. S2 E). Taken together, these data show that SEPT9 interacts directly and specifically with dynein-dynactin and alone or in complex with SEPT2/6/7 triggers retrograde trafficking by recruiting the dynein motor.

### SEPT9 interacts with both dynein and dynactin through its GTP-binding and N-terminal extension domains, respectively

Activation of dynein-driven motility requires adaptor proteins that promote the assembly and stability of dynein-dynactin complexes. Activating dynein adaptors are characterized by long coiled-coil domains, which bridge dynein with dynactin by interacting concomitantly with subunits of the dynein-dynactin complex (Cross and Dodding, 2019; McKenney et al., 2014; Reck-Peterson et al., 2018). Thus, we sought to determine how SEPT9 interacts with dynein-dynactin.

SEPT9 consists of an N-terminal extension (NTE) domain, which is largely disordered (Bai et al., 2016), and a GTP-binding domain (G-domain) that is evolutionarily and structurally related to small GTPases. We purified recombinant versions of these domains and tested whether they interact directly with native dynein and dynactin. Strikingly, we found that the NTE and G-domain have differential affinities for dynein and dynactin. The dynactin subunit p150^Glued^ was pulled down robustly with the NTE domain, but it was barely detectable in pull-downs with the G-domain of SEPT9 (Fig. 3 A). Conversely, the G-domain associated strongly with DHC, which exhibited weak binding to the NTE domain (Fig. 3 A). Hence, SEPT9 can interact with both dynein and dynactin through its GTP-binding and NTE domains, respectively.

**Figure 3.**
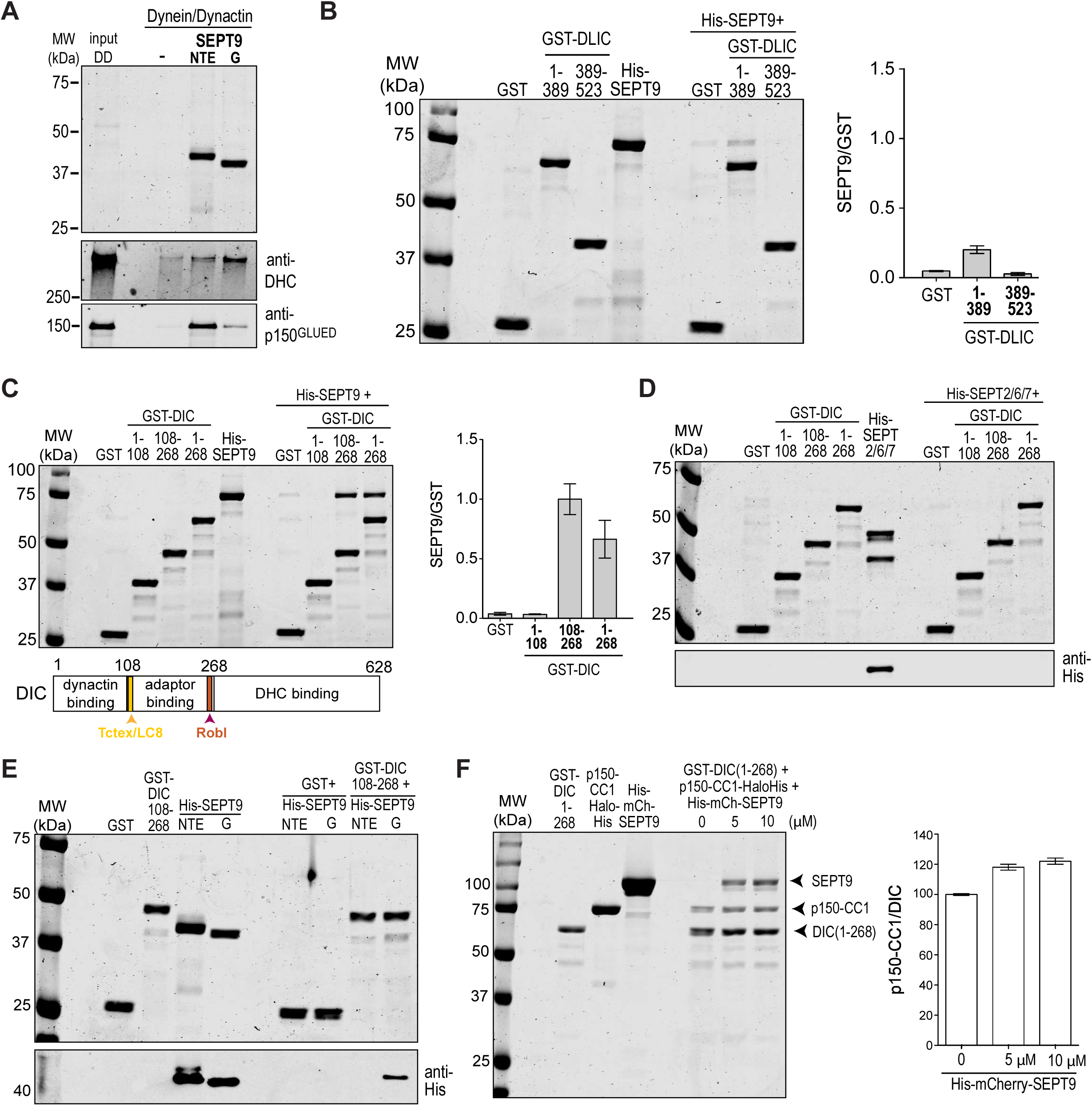
SEPT9 provides a molecular link between dynein and dynactin through its GTP-binding and NTE domains, respectively. (A) Coomassie-stained gel shows the recombinant NTE (N) and GTP-binding (G) domains of SEPT9, which were used in pull-down assays with purified dynein and dynactin. Pull-downs were western blotted with antibodies against DIC and p150^GLUED^. (B) Coomassie-stained gel shows the results of protein binding assays between recombinant His-SEPT9 and GST (control) or the GST-tagged N-(aa 1-389) and C-terminal (aa 389-523) halves of DLIC. Bar graph shows the mean ratio of SEPT9 to GST or GST-DLIC protein band intensities from three independent experiments; error bars indicate S.E.M. (C) Coomassie-stained gel shows results from pull-down assays of His-SEPT9 with GST (control) or GST-tagged DIC(1-108), DIC (108-268) and DIC(1-268). Schematic depicts the major domains of DIC and their corresponding interactions with components of the dynein-dynactin complex. Bar graph shows quantification of SEPT9 pull-down as the mean ± SEM ratio value of SEPT9 to GST protein band intensities from five independent experiments. (D) Coomassie-stained gel (top) and western blot (bottom; anti-His) show results from pull-down assays of purified recombinant His-SEPT2/6/7 with GST (control) or GST-tagged DIC(1-108), DIC(108-268) and DIC(1-268). (E) Coomassie-stained gel (top) and western blot (bottom; anti-His) show results from pull-down assays of recombinant NTE and G domain fragments of His-SEPT9 with GST-DIC(108-268). (F) Gel shows the input and results from protein binding assays of equimolar (0.5 μM) recombinant p150^GLUED^-CC1-Halo-His with GST-DIC(1-268) in the presence of increasing concentrations of His-mCherry-SEPT9 (0, 5, 10 μM). Bar graph shows the relative increase in the amount of DIC(108-268)-bound p150^GLUED^-CC1 with increasing concentrations of SEPT9; ratio was set to 100 for 0 μM of SEPT9. Error bars show median, low and high values of protein band intensity ratios from three independent experiments.

To gain more insight into the SEPT9-dynein interactions, we examined if SEPT9 associates with domains of DLIC, which are critical for the activation of dynein motility by interacting directly with dynein adaptors. We tested if SEPT9 interacts with the C-terminus (389-523 aa) of DLIC, which is a common binding site for several dynein activators (Gama et al., 2017; Lee et al., 2018; Schroeder et al., 2014; Schroeder and Vale, 2016). SEPT9 did not bind DLIC(389-523) and similarly had a weak affinity for the N-terminal domain of DLIC (Fig. 3 B). In contrast, SEPT9 exhibited a robust interaction with the N-terminal domain of DIC (1-268 aa), which has been shown to interact with the snapin and huntingtin adaptors (Caviston et al., 2007; Di Giovanni and Sheng, 2015). While SEPT9 did not interact with the first 108 N-terminal amino acids of DIC, which associate with the p150^GLUED^ subunit of dynactin, it bound DIC(108-268), which interacts with dynein light chains and the dynein adaptor snapin (Fig. 3 C) (Di Giovanni and Sheng, 2015). Full length SEPT9 and its G-domain interacted specifically with DIC(108-268), while neither the NTE domain of SEPT9 or SEPT2/6/7 associated with DIC(108-268) (Fig. 3 D-E). These data indicate that SEPT9-dynein binding involves a direct interaction between the G-domain of SEPT9 and DIC(108-268).

Given that the interaction of DIC with p150^GLUED^ is critical for the modulation of dynein motility (Ayloo et al., 2014; Culver-Hanlon et al., 2006; Feng et al., 2020; King and Schroer, 2000; Vaughan and Vallee, 1995), we tested whether SEPT9 impacts the association of the CC1 domain of p150^GLUED^ with the N-terminus of DIC. Using purified proteins, we assayed for direct binding of DIC(1-268) to p150^GLUED^-CC1 in the presence of increasing concentrations of recombinant SEPT9. SEPT9 enhanced binding of p150^GLUED^-CC1 to the N-terminal fragment of DIC by ∼20% (Fig. 3 F). Albeit modest, this was a reproducible effect with two different concentrations of SEPT9, suggesting that SEPT9 impacts DIC-p150^GLUED^ binding. Thus, similar to *bona fide* adaptors of dynein, SEPT9 interacts concomitantly with dynein and dynactin and may facilitate cooperative assembly into processive motor complexes.

### SEPT9 functions as a GDP-activated switch of dynein-driven motility

Septins are a large family of paralogs, which vary in their ability to hydrolyze GTP (Sirajuddin et al., 2009; Zent et al., 2011; Zent and Wittinghofer, 2014). GTP-hydrolyzing septin paralogs such as SEPT9 assemble into functional oligomers and multimers that are bound to GDP (Castro-Castro et al., 2016). In contrast to monomeric small GTPases, which are active in their GTP-bound state, GTP-hydrolyzing septins are functionally active as GDP-bound multimers.

Association of SEPT9 with DIC through its G-domain raised the possibility that SEPT9 interacts with dynein and induces dynein-driven transport in a nucleotide-dependent manner. To examine whether the G-domain of SEPT9 interacts preferentially with DIC in its GTP- or GDP-bound state, we performed protein-binding assays with recombinant SEPT9 G-domain and DIC(108-268) in the presence of GDP, GTP and non-hydrolyzable GTPγS. Strikingly, both GTP and non-hydrolyzable GTPγS abrogated SEPT9-DIC binding, which was observed only in the presence of GDP (Fig. 4 A). To further test whether SEPT9 binds DIC in its GDP-bound state, we mutated the catalytic threonine (T339) of the G-domain into a glycine, a residue that is conserved in place of the catalytic threonine in septin paralogs that are constitutively bound to GTP (Sirajuddin et al., 2009). Compared to wild-type SEPT9 G-domain, binding of the T339G mutant to DIC(108-268) was markedly reduced (Fig. 4 B). Thus, GTP hydrolysis is critical for the interaction of SEPT9 with dynein.

**Figure 4.**
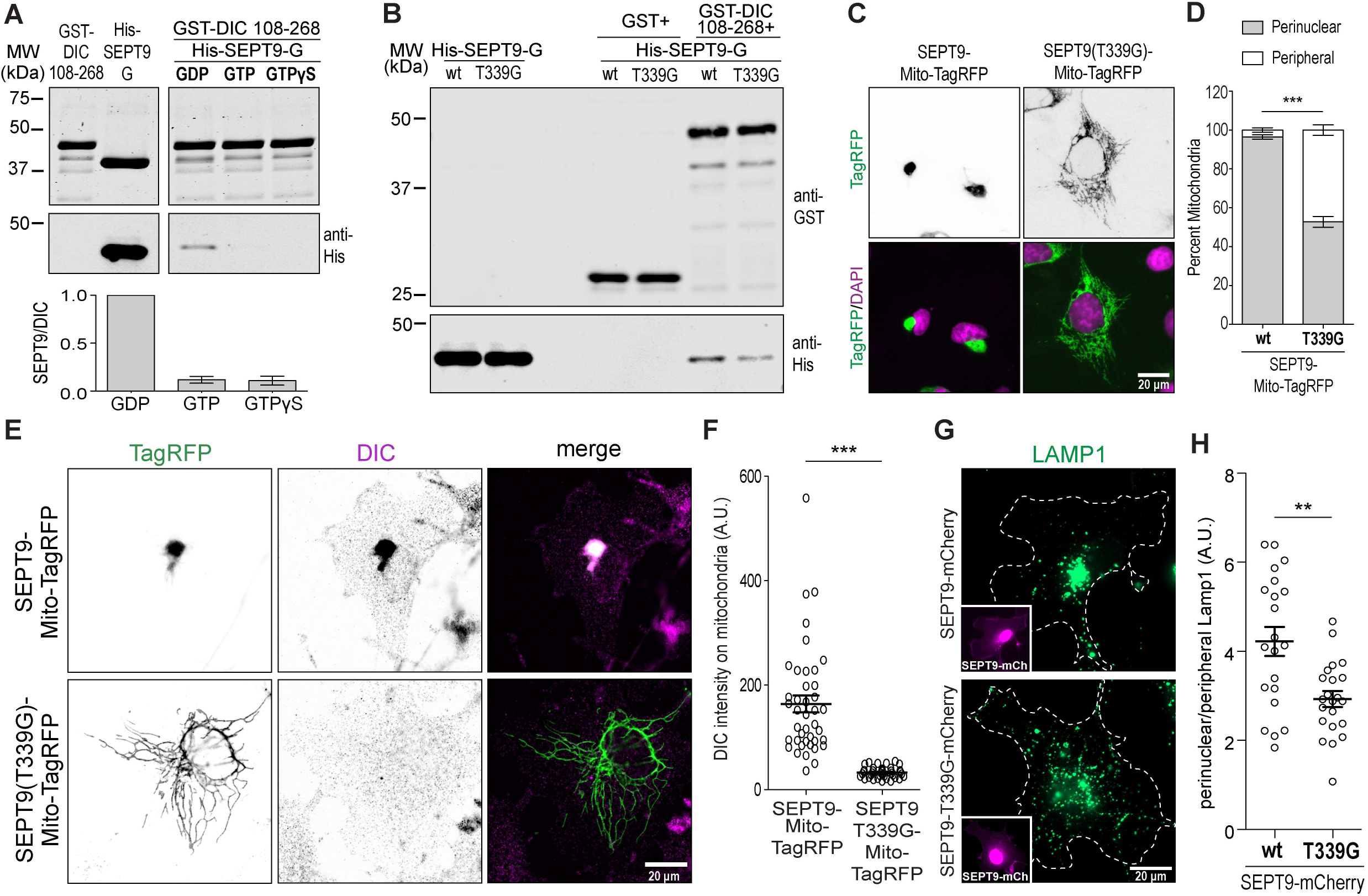
The GDP-bound state of SEPT9 recruits dynein and induces retrograde transport. (A) Coomassie-stained gel (top) and western blot (bottom; anti-His) show binding of the G-domain of SEPT9 to GST-DIC(108-268) in the presence of GDP, GTP or GTPγS (0.5 mM). Quantification shows the ratios of SEPT9 G-domain to GST-DIC(108-268) protein band intensities from three independent experiments. (B) Gels show western blots (anti-GST/anti-His) of pull-downs of recombinant His-tagged wild-type or GTPase-dead (T339G) SEPT9 with GST or GST-DIC(108-268). (C) Images show DAPI-stained COS-7 cells transfected with mitochondria-targeted (Mito-TagRFP) wild-type and GTPase-dead (T339G) SEPT9. (D) Bar graph shows perinuclear and peripheral mitochondria as percentage (mean ± SEM) of total mitochondria per cell (*n* = 24-27). (E) Images show COS-7 cells transfected with mitochondria-targeted (MitoTagRFP) wild-type or GTPase-dead (T339G) SEPT9 and stained with antibodies against DIC. (F) Plot shows DIC fluorescence intensity (mean ± SEM) on mitochondria with MitoTagRFP-SEPT9 or SEPT9(T339G)-Mito-TagRFP per cell (*n* = 40-41). (G) Images show the distribution of lysosomes (LAMP-1) in COS-7 cells transfected with wild type SEPT9 or SEPT9(T339G) (insets). (H) Plot shows the ratio (mean ± SEM) of perinuclear to peripheral LAMP1 fluorescence intensity per cell (*n* = 21-22). **, p < 0.01; *** p < 0.001.

Next, we examined if the dynein-driven motility of membrane organelles is similarly sensitive to the nucleotide-bound state of SEPT9. Targeting of SEPT9(T339G) to mitochondria did not induce the perinuclear clustering observed with wild type SEPT9 (Fig. 4C-D). Moreover, SEPT9(T339G) failed to recruit DIC to mitochondria, demonstrating that membrane recruitment of DIC requires GDP-bound SEPT9 (Fig. 4 E-F). Over-expression of SEPT9(T339G) did not impact the intracellular position of lysosomes, which did not accumulate to the perinuclear cytoplasm as observed with SEPT9 (Fig. 4 G-H). Collectively, these data show that SEPT9 recruits dynein and induces dynein-driven motility in its GDP-bound state, which favors SEPT9 dimerization and assembly into higher order multimers. Thus, SEPT9 provides a GDP-activated scaffold for the recruitment and assembly of dynein-dynactin motors.

### SEPT9 promotes dynein-driven transport of lysosomes in response to acute oxidative stress

Our results suggest that SEPT9 provides a novel and alternative mechanism to the canonical small GTPase-based recruitment of dynein to lysosomes. In contrast to Rab7, which requires GTP, dynein recruitment by SEPT9 is GDP-activated. We posited that this novel septin GTPase mechanism could be utilized in adaptive cellular responses to stress.

To investigate if SEPT9 is involved in the retrograde trafficking of lysosomes under conditions of cell stress, we induced acute oxidative stress with sodium arsenite (NaAsO_2_), which triggers release of reactive oxygen species (ROS) (Ellinsworth, 2015). After 30 minutes of arsenite treatment, there was a striking loss of subnuclear septin filaments and an amplification of perinuclear SEPT9 puncta (Fig. 5 A). Quantitatively, SEPT9 localization to lysosomes (LAMTOR4) was markedly enhanced in arsenite-treated cells (Fig. 5 B). Under conditions of glucose deprivation, which also results in ROS production and oxidative stress (Graham et al., 2012; Liu et al., 2003; Song and Hwang, 2018), SEPT9 filaments were not disrupted and lysosomal levels of SEPT9 puncta did not increase (Fig. 5 B). Thus, enhancement of SEPT9 localization to lysosomes appears to occur specifically in response to arsenite-induced stress.

**Figure 5.**
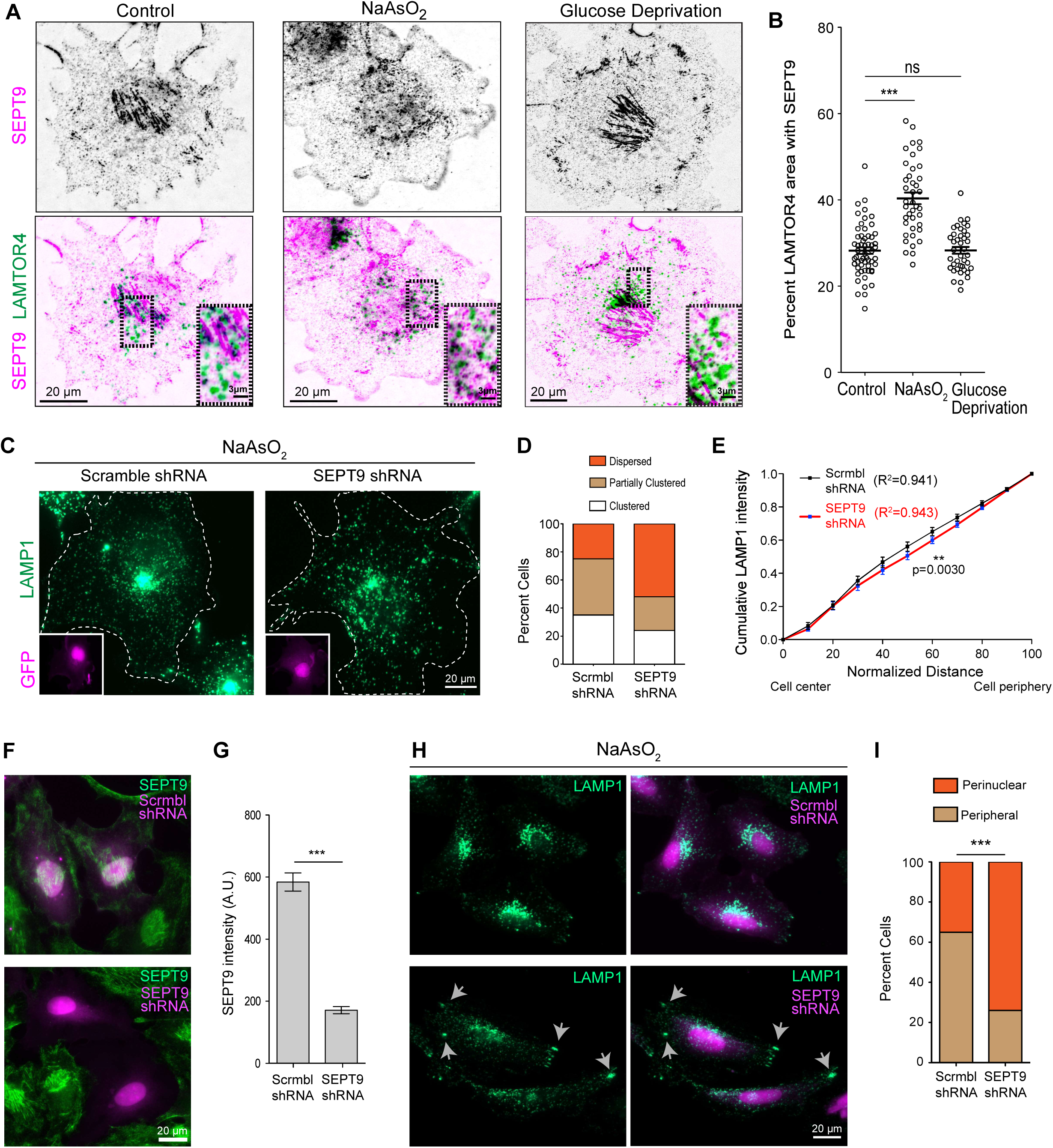
Oxidative stress increases lysosomal levels of SEPT9, which promotes perinuclear clustering. (A) Confocal microscopy images (inverted monochrome) of COS-7 cells treated with NaAsO_2_ (300 μM) or deprived of glucose for 1 h, and stained for LAMTOR4 (green) and SEPT9 (magenta). Outlined perinuclear regions are shown in higher magnification. (B) Bar graph shows the percentage (mean ± SEM) of total lysosomal area (LAMTOR4) that overlaps with endogenous SEPT9 per cell (*n* = 40-59) after NaAsO_2_ (300 μM, 30 minutes) treatment or glucose deprivation (1 h). (C) COS-7 cells were transfected with GFP-expressing scramble or SEPT9 shRNAs (insets) and treated with NaAsO_2_ (300 μM) for 2 h. Images show lysosome (LAMP-1) localization. (D) Bar graph shows percentage of cells (*n* = 20-21) with dispersed, perinuclearly clustered and partially clustered lysosome distribution. (E) Plot shows the fraction of total LAMP-1 intensity per COS-7 cell area normalized (percent of total) to distance from cell center to peripheral edge in control and NaAsO_2_ conditions (*n* = 15-18). (F) HeLa cells were transfected with GFP-expressing scramble or SEPT9 shRNAs (magenta), and stained for SEPT9 (green). (G) Bar graph shows mean (± SEM; *n* = 65-66) fluorescence intensity of SEPT9 per shRNA-expressing cell. (H) HeLa cells were transfected with GFP-expressing scramble or SEPT9 shRNAs (magenta), and treated with NaAsO_2_ (400 μM) for 1 h. Images show LAMP1 distribution and arrows point to clusters of lysosomes at peripheral cell edges. (I) Bar graph shows percentage of cells with peripheral or perinuclear without any peripheral clusters of lysosomes (n = 99-115). Ns, non-significant (p > 0.05); *, p < 0.05; **, p < 0.01; *** p < 0.001.

To determine whether SEPT9 contributes to the dynein-driven transport of lysosomes under oxidative stress, we analyzed the intracellular distribution of lysosomes in control and SEPT9-depleted cells after a two-hour treatment with sodium arsenite, which causes lysosomal congresion to the perinuclear cytoplasm (Willett et al., 2017). SEPT9 knock-down increased the percentage of COS-7 cells with a dispersed lysosomal distribution (Fig. 5 C-D), and lysosome presence was reduced in cytoplasmic areas of an intermediate distance from the center of the cell to the peripheral edge (Fig. 5 E). In HeLa cells, SEPT9 depletion had a similar effect on the perinuclear accumulation of lysosomes, which was markedly reduced after one hour of arsenite treatment (Fig. 5 F-I). Moreover, clusters of lysosomes localized to peripheral areas of the cell edge as if they were unable to move retrogradely toward the nucleus (Fig. 5H, arrows). This phenotype is reminiscent of the peripheral accumulation of lysosomes in cells depleted of the dynein adaptor JIP-4 (Willett et al., 2017). Taken together, these data demonstrate that SEPT9 is critical for the retrograde transport of lysosomes under conditions of oxidative stress.

## Discussion

The intracellular traffic and position of lysosomes are critical for their biogenesis, turnover and physiological functions. In response to a variety of metabolic and signaling cues, lysosomes mobilize by recruiting kinesin and/or dynein motors, but the underlying mechanisms are not well understood. Here, we have discovered a novel mechanism of dynein-driven motility, which is mediated by SEPT9, a GTPase of the septin family. Unlike the Rab7 GTPase and lysosomal membrane proteins, which recruit dynein through cytoplasmic adaptor proteins (e.g., RIPL, JIP4, ALG4), SEPT9 is a membrane-associated GTPase that interacts directly with dynein. Notably, SEPT9 associates preferentially with dynein in its GDP-bound state, which favors SEPT9 dimerization and assembly into higher order oligomers. Hence, in contrast to the monomeric small GTPases of the Rab and Arf families, which are activated by GTP, SEPT9 provides a GDP-activated scaffold for the recruitment of multiple dynein-dynactin complexes. This scaffolding function may facilitate cooperative assembly of dynein-dynactin into the nanodomains of motor teams, which have been observed on lipid microdomains and microtubules (Cella Zanacchi et al., 2019; Rai et al., 2016)

Among the members of the septin family of GTPases, SEPT9 is unique in possessing a faster GTPase activity and a dimerization interface, which is characterized by nucleotide-dependent plasticity (Castro et al., 2020; Zent and Wittinghofer, 2014). Additionally, SEPT9 has been reported to function and localize independently of its heteromeric partners as the central subunit of the octameric SEPT2/6/7/9 complex (Estey et al., 2010; Karasmanis et al., 2018). Our results indicate that SEPT9 interacts with dynein independently of SEPT2/6/7, but do not rule out the possibility that SEPT9 functions in a complex with SEPT2/6/7 on lysosomal membranes. The faster hydrolytic activity of SEPT9 may select for its homomeric assembly and concomitantly favor dynein-binding. Recent crystallographic studies show that GDP shifts the dimeric interface between the N- and C-termini of SEPT9 into a distinct open conformation, which has not been observed in other septin paralogs (Castro et al., 2020). This shift from a closed GTP-bound to an open GDP-bound interface impacts differentially the two polybasic membrane-binding domains of SEPT9, one of which becomes occluded while the other disengages from a neutralizing polyacidic domain (Castro et al., 2020; Omrane et al., 2019). Therefore, the GDP-bound SEPT9 could interact with dynein while in a membrane-binding mode that favors association with lysosomes.

Similar to dimeric *bona fide* dynein adaptors, which activate the processive motility of dynein, SEPT9 interacts concomitantly with dynein and dynactin. However, SEPT9 lacks the DLIC-binding domains and coiled coil sequences of activating adaptors, and we were not able to observe an enhancement of dynein processivity in single molecule *in vitro* motility assays. If SEPT9 alone is not sufficient to activate dynein-dynactin motility, we surmise that it provides a scaffold for the assembly of multiple dynein complexes, whose activation depends on the recruitment of additional factors (Rai et al., 2016; Urnavicius et al., 2018). Additionally, SEPT9 may reinforce dynein tethering to the membrane cargo and/or microtubules, facilitating movement under high load. As a dynein scaffold, SEPT9 could also coordinate dynein movement with kinesin motors. Notably, SEPT9 interacts with the cargo-binding C-terminal tail of kinesin-2/KIF17 (Bai et al., 2016) and therefore may provide a mechanism for switching between microtubule motors of opposite directionality as previously shown for the JNK-interacting protein-1 (JIP-1) (Fu and Holzbaur, 2013). Coincidently, SEPT9 also interacts with JNK, which triggers a dynein-kinesin switch by phosphorylating JIP-1 (Gonzalez et al., 2009).

Previous studies have implicated septins in endocytic trafficking to lysosomes and autophagic-lysosomal delivery, both of which involve dynein-mediated transport (Song et al., 2016). Septins are critical for the biogenesis of multivesicular bodies (Traikov et al., 2014), and SEPT9 associates with the tumor susceptibility gene 101 (TSG101), a component of the endosomal sorting complex required for transport (ESCRT) that binds ubiquitinated cargo and receptors (Karasmanis et al., 2019). It is plausible that SEPT9 links TSG101 to dynein, selecting for the retrograde flux of endosomes and multivesicular bodies (MVBs) with ubiquitinated proteins toward lysosomes. The dynein adaptor RILP has been similarly observed to interact with the ESCRT-II subunits Vps22 and Vps36, and is required for the biogenesis of multivesicular endosomes and the degradation of epidermal growth factor receptors (Progida et al., 2007; Progida et al., 2006). A septin-mediated retrograde transport of lysosomes and autophagosomes may also promote the autophagic destruction of pathogenic bacteria such as *Shigella flexneri*, which are entrapped in septin cage-like structures (Torraca and Mostowy, 2016). Septins are required for the recruitment of autophagic components to encaged bacteria and subsequent merging with lysosomes (Krokowski et al., 2018; Mostowy et al., 2010), which might be facilitated by retrograde movement to the cell center.

Considering that SEPT9 associates with microtubules, our findings pose the question of whether and how SEPT9 functions in dynein-driven traffic as both a microtubule-associated protein and membrane scaffold/adaptor. Results from *in vitro* motility assays have indicated that microtubule-associated SEPT9 inhibits dynein motility (Karasmanis et al., 2018), while membrane-associated SEPT9 promotes dynein-mediated transport. While these roles are not mutually exclusive, SEPT9 could shift from microtubules to endomembranes in response to cellular conditions and signaling cues. In serum-starved cells, loss of filamentous cytoskeleton-associated septins has been reported previously (Kinoshita et al., 1997), and our data show that SEPT9 localization to lysosomes is enhanced under cell stress. The N-terminal domain of SEPT9, which interacts with microtubules and actin filaments, contains a number of putative phosphorylation sites, indicating that SEPT9 localization is under phospho-regulation by signaling kinases. Hence, an on-demand shift of SEPT9 from microtubules to lysosomes would enable dynein-driven movement without interference from microtubule-associated SEPT9 molecules. Interestingly, this shift might be a property that characterizes a number of microtubule-associated proteins including the dynein adaptor HOOK-1 which associate with both microtubules and membrane organelles (Krtkova et al., 2016; Linden et al., 1989; Maldonado-Baez et al., 2013; Tortosa et al., 2017).

In sum, our findings bear broader significance for dynein-driven transport and cellular adaptation to stress. Beyond lysosomes, septins can function as membrane scaffolds for the assembly and activation of dynein-dynactin complexes on a variety of endomembrane organelles and plasma membrane domains of micron-scale curvature, which favors membrane-septin binding. For example, septins could recruit dynein-dynactin on organelles such as lipid droplets, whose growth and perinuclear accumulation depends on SEPT9 during hepatitis C virus infection (Akil et al., 2016). Alternatively, septins can facilitate dynein-mediated capture and tethering of microtubules at cortical membrane sites and possibly enable dynein-mediated microtubule focusing and sliding (Hendricks et al., 2012; Ligon et al., 2001; Tanenbaum et al., 2013). Importantly, under conditions of GTP shortage or low GTP/GDP ratios, septins could provide a mechanism of dynein-dynactin motility, which can be activated by GDP rather than GTP as needed for the small GTPases such as Rab7. Recent findings of activation of the mammalian target of rapamycin complex 2 (mTORC2) by Rho-GDP suggests that GTPase modules can adopt GDP-activated functions (Senoo et al., 2019). Thus, a GDP-triggered assembly of septins into higher order membrane scaffolds might serve as an adaptive and/or energy-sensing mechanism for the dynein-driven transport of lysosomes under conditions of cell stress.

## Materials and Methods

### Antibodies and reagents

Cells were immunostained with the following antibodies: mouse anti-Lamp1 (1:100; Iowa DSHB), rabbit anti-SEPT9 (1:100; Proteintech Group), rabbit anti-SEPT9 (1:200; Novus Biologicals), mouse anti-SEPT9 clone 10C10 (1:100; Millipore-SIGMA), rabbit anti-SEPT2 (a5N5 1:300; gift from Makoto Kinoshita, Nagoya University, Japan), rabbit anti-SEPT7 (1:300; IBL America), rabbit anti-SEPT6 (1:300; gift from Makoto Kinoshita, Nagoya University, Japan), mouse anti-GM130 (1:200; BD Biosciences), mouse anti-DIC antibody clone 74.1 (1:100; Millipore), rabbit anti-LAMTOR4 (1:200; Cell Signaling), mouse anti-EEA1 clone 14/EEA1 (1:100; BD Biosciences), anti-alpha/beta tubulin (1:100; Cytoskeleton). F(ab’)_2_ fragment affinity-purified secondary antibodies (1:200) were purchased from Jackson ImmunoResearch Laboratories and included donkey anti-mouse, -rabbit, -chicken and -goat antibodies conjugated with AMCA, Alexa488, Alexa594 or Alexa647. Cells were counterstained with DAPI (SIGMA), Phalloidin CF405 (1:200; Biotium) or MitoSpy™ NIR DiIC1 dye (BioLegend) as indicated and according to the manufacturer’s instructions. Western blots were performed with mouse anti-Lamp1 (1:500; Iowa DSHB), rabbit anti-SEPT9 (1:1000; Proteintech Group), rabbit anti-SEPT2 (a5N5 1:2000; gift from Makoto Kinoshita, Nagoya University, Japan), rabbit anti-SEPT7 (1:2000; IBL America), rabbit anti-SEPT6 (1:1000; gift from Makoto Kinoshita, Nagoya University, Japan), mouse anti-GM130 (1:1000; BD Biosciences), mouse anti-Rab7A (1:500; Abcam), rabbit anti-Rab5 (1:500; Santa Cruz), anti-Bip, mouse anti-DIC antibody clone 74.1 (1:1000; Millipore), mouse anti-p150^Glued^ (1:1000; BD Biosciences), mouse anti-mCherry (1:2000; Abcam), rabbit anti-DynC1H1 (1:500; Proteinteck Group), mouse anti-His tag (1:2000; QIAGEN) and rabbit anti-GST (1:2000; Santa Cruz). Secondary antibodies for Western blots (1:10,000) were purchased from LI-COR including donkey anti-mouse and anti-rabbit 800CW or 680RD conjugates. Nucleotides were purchased from SIGMA (GTPγS, GDP) and Cytoskeleton, Inc (GTP).

### Plasmids and constructs

Plasmids encoding for GFP-EEA1 was a gift from Silvia Corvera (Addgene plasmid # 42307; http://n2t.net/addgene:42307; RRID:Addgene_42307) (Lawe et al., 2000), LAMP1-mGFP was a gift from Esteban Dell’Angelica (Addgene plasmid #34831; http://n2t.net/addgene:34831; RRID:Addgene_34831) (Falcon-Perez et al., 2005), pGEX6P1-human LIC1 G domain (GST-LIC 1-389) was a gift from Ron Vale (Addgene plasmid # 74598; http://n2t.net/addgene:74598; RRID:Addgene_74598) (Schroeder et al., 2014), pGEX6P1-human LIC1 C-terminal half (GST-LIC 389-523) was a gift from Ron Vale (Addgene plasmid #74599, http://n2t.net/addgene:74599; RRID:Addgene_74599) (Schroeder et al., 2014) and EGFP-Rab7A T22N was a gift from Qing Zhong (Addgene plasmid # 28048; http://n2t.net/addgene:28048; RRID:Addgene_28048) (Sun et al., 2010). The plasmid for PEX-RFP-FKBP (Kapitein et al., 2010) was a gift from Casper Hoogenraad (Utrecht University), GFP-p50dynamitin (Quintyne et al., 1999) was a gift from Trina Schroer (John Hopkins University) and MitoTagRFP (N1-TagRFP-ActA) (Sirianni et al., 2016) was donated by Adam Kwiatkowski (University of Pittsburgh). Plasmids expressing recombinant GST-DIC 108-268 and GST-DIC 1-268 were kindly donated by Dr. Zu-Hang Sheng (National Institutes of Health) (Di Giovanni and Sheng, 2015), p150^GLUED^-CC1-Halo-6xHis (216-547aa) was a kind gift from Dr. Erika Holzbaur (University of Pennsylvania) (Ayloo et al., 2014)and sfGFP-BICD2 (25-400aa) was provided by Dr. Richard McKenney (University of California, Davis) (McKenney et al., 2014).

The plasmids GFP-Rab5A (Dolat and Spiliotis, 2016), mCherry-N1-humanSEPT9_i1 (Bai et al., 2013), mCherry-N1-SEPT2 (Bowen et al., 2011), pmCherry-N1-ratSEPT9 and pET28a-mCherry-humanSEPT9 (Karasmanis et al., 2018), pET28a-SEPT9(i1), pET28a-SEPT9-NTE (1-283 aa) and pET28a-SEPT9G (283-586 aa) (Bai et al., 2016) were constructed as previously described. The pnEA-vH and pnCS vectors encoding for His-SEPT2 and SEPT6/7-strep, respectively, have been described previously (Nakos et al., 2019).

Plasmids encoding for mitochondria-targeted septins were made as follows: mouse SEPT2, human SEPT6 and rat SEPT7 were PCR amplified using the primers 5’-CAT CTC GAG ATG TCT AAG CAA CAA C-3’ and 5’-ACT GGA TCC CAC ACA TGC TGC CCG AG-3’, 5’-CAT CTC GAG ATG GCA GCG ACC GAT ATA G-3’ and 5’-ACT GGA TCC CAA TTT TTC TTC TCT TTG TC-3’, 5’-CAT CTC GAG ATG TCG GTC AGT GCG-3’ and 5’-ACT GGA TCC CAA AAG ATC TTG CCT TTC-3’, respectively and inserted in the pTagRFP-Mito plasmid using XhoI, BamHI sites. Human SEPT9_i1 (NM_001113491.2) was PCR amplified using the primers 5’-CAT CTC GAG ATG AAG AAG TCT TAC-’3, 5’-ACT AAG CTT CAT CTC TGG GGC TTC-3) and inserted into pTagRFP-Mito using XhoI, HindII sites. pmCherry-C1-SEPT6 was contructed by PCR amplification of human SEPT6 using the primers 5’-TCG AAG CTT CCA TGG CAG CGA CCG ATA TAG-3’, 5’-TCG GGA TCC TTA ATT TTT CTT CTC-3’ and subsequent cloned into pmCherry-C1 with HindIII/BamHI sites. To construct pmCherry-SEPT9-PGK, SEPT9-mCherry (NM_001113491.2) was subcloned from pmCherry-N1-SEPT9 into the NheI & NotI sites of the pcDNA-PGK-MCS-Hyg (gift from Dr. Kay Oliver Schink, Norwegian Radium Hospital).

pEGFP-C2-SEPT9-FRB was constructed by inserting FRB sequence, amplified from GFP-BICDN-FRB (gift from Casper Hoogenraad, Utrecht University) (Hoogenraad et al., 2003), into pEGFP-C2 vector using EcoRI and BamHI. SEPT9 (NM_001113491.2) was subsequently cloned into the pEGFP-C2-FRB plasmid with XhoI/HindIII using the following primers; 5’-AAAAAACTCGAGCATGAAGAAGTCTTACTCA-3’ and 5’-TTTTTTAAGCTTGACATCTCTGGGGCTT-3’. pGEX-KT-ext-DIC 1-108aa (mouse IC1A) was created by inserting the PCR-amplified fragment of DIC using the primers 5’-AAAAAAGGATCCATGTCTGACAAGAGCGAC-3’, 5’-TTTTTTCTCGAGCTACTGCAGGGTCCT-3’ with BamH1/XhoI. For constructing targeting SEPT9 shRNA (5’-G CAC GAT ATT GAG GAG AAA-3’) against human and monkey SEPT9 and scrambled non-targeting shRNA control (5-G CGA AGA AGG ATA CGT AAT-3), targeting sequences were inserted into the pSuper-GFP vector. Targeting sequence of SEPT9 shRNA was designed using BLOCK-iT™ RNAi Designer (Thermofisher) and the sequence was scrambled using the Invivogen shRNA wizard.

pET-28a-SEPT9-Gdomain-T339G, pTagRFP-Mito-SEPT9-T339G and pmCherry-SEPT9-T339G mutants were made according to KAPA Biosystems Site-directed mutagenesis protocol. Briefly, the above-mentioned SEPT9 plasmids were amplified with KAPA HiFi HotStart DNA polymerase (KAPA BIOSYSTEMS) using the primers 5’-GGAGCGCATCCCCAAGGGCATCGAGATCAAGTCC-3’ and 5’-GGACTTGATCTCGATGCCCTTGGGGATGCGCTCC-3’. The amplified PCR product was treated with DpnI (New England Biolabs) for 1 h at 37 °C, heat inactivated at 80°C for 20 min and was subsequently transformed in *E. coli* DH5a competent cells.

### Expression and purification of recombinant proteins

Plasmids encoding for recombinant His-tagged proteins were transformed into *Escherichia coli* BL21(DE3) (Invitrogen). Bacterial cultures were grown at 37°C in LB media to OD_600_ of 0.6-0.8 and induced with 0.5 mM IPTG at 18 °C for 16h (overnight). Cells were harvested by centrifugation at 4,000rpm in a JA-10 fixed angle rotor (Beckman Coulter). Pellets were resuspended in lysis buffer containing 50 mM Tris, pH 8, 300 mM NaCl, 10% glycerol, and 10 mM imidazole supplemented with 2mM PMSF (Sigma), a Protease inhibitor cocktail (G-Biosciences) and 10µg/ml DNAse I (Millipore). Bacteria were lysed by sonication and clarified by centrifugation at 20,000 xg for 30 min in a JA-20 rotor (Beckman) at 4°C. Supernatant was loaded onto 1 ml Ni-NTA agarose beads (Macherey-Nagel), pre-equilibrated with 10 ml wash buffer and rotated end-over-end for 1h at 4°C. Beads were washed with 30 ml wash buffer (50 mM Tris pH 8.0, 300 mM NaCl, 10% glycerol, 10 mM imidazole, 2mM PMSF) in a gravity flow column and protein was eluted with elution buffer containing 300 mM imidazole (50 mM Tris pH 8.0, 300 mM NaCl, 10% glycerol, 2mM PMSF and 300 mM imidazole). Peak fractions were combined and dialyzed overnight against 50 mM Tris, pH 8.0, 150 mM NaCl, and 10% glycerol. For proteins used in single motility assays (His-mCherry-SEPT9) and p150^Glued^ CC1-Halo-6xHis, peak fractions eluted from Ni-NTA beads were further purified via size exclusion chromatography on a Superdex 200 (GE Healthcare Life Sciences) with degassed dialysis buffer (50 mM Tris, pH 8.0, 150 mM NaCl, and 10% glycerol) at 0.5 ml/min. Peak fractions were collected and dialyzed against GF150 buffer (25 mM HEPES pH7.4, 150 mM KCl, 1mM MgCl_2_, 1 mM DTT, 10% glycerol) overnight (Schlager et al., 2014). Proteins were aliquoted, snap frozen and stored at -80 °C. GST tagged proteins were purified with the following modifications. Bacteria pellets were lysed in PBS containing 10% glycerol and supplemented with 2mM PMSF (Sigma), a Protease inhibitor cocktail (G-Biosciences) and 10µg/ml DNAse I (Millipore). Cleared lysates were loaded onto GST-Trap (GE Healthcare Life Sciences) washed with 20-30 column volumes wash buffer (PBS, 10% glycerol and 2mM PMSF) at 0.5 ml/min and eluted with 50mM glutathione (SIGMA) (wash buffer supplemented with 50mM glutathione). Peak fractions were dialyzed overnight against dialysis buffer (50 mM Tris, pH 8.0, 150 mM NaCl, and 10% glycerol). SEPT2/6/7 complex was expressed and purified as previously described (Nakos et al., 2019) with the modification of dialyzing the peak fraction elution from the StrepTrap HP column in buffer consisting of 50 mM Tris, pH 8.0, 150 mM NaCl and 10% glycerol.

### Dynein-dynactin purification

Native dynein/dynactin was purified from HEK-293T cells as previously described (Huynh and Vale, 2017; McKenney et al., 2014). Briefly, ten to fifteen plates (15 cm) were harvested per purification. Cells were washed twice with PBS and collected by scraping with a flexible blade cell scraper (VWR). Cells were pelleted at 1000 x g for 5 minutes and lysed in 2 cell-pellet volumes of Buffer-A (30 mM Hepes, pH 7.4, 50 mM potassium acetate, 2 mM magnesium acetate, 1 mM EGTA, and 10% glycerol) supplemented with 2 mM PMSF, 5 mM DTT, 0.2% NP-40 + 0.1 mM ATP and a protease inhibitor cocktail (SIGMA). Lysate was rotated end-over-end for 1h at 4°C to ensure lysis. Lysate was clarified at 50K for 30 min in a TLA 100.3 type rotor (Beckman Coulter) at 4°C. The clarified supernatant was mixed with 200nM sfGFP-BICDN and 50 µl Streptactin beads (IBA Life Sciences) and rotated end-over-end overnight at 4 °C. Beads were pelleted by centrifugation at 500 xg for 1 minute and washed five times with 500 µl of Buffer A supplemented with 2mM PMSF, 5mM DTT, 0.1% NP-40. Dynein and dynactin were eluted from sfGFP-BICDN beads by incubating for 15 min with 50-100 μl buffer A containing 300 mM NaCl and 0.5 mM ATP on ice. The eluate was passed through a spin filter to remove any remaining beads. Eluate was diluted with two volumes of Buffer-A to reduce salt concentration to 100 mM and 6% sucrose was added before aliquoting and snap freezing.

N-terminally Strep-II sfGFP BICD2 (25-400aa) (gift from Richard McKenney, University of California, Davis) was expressed in BL21-DE3 cells and purified as mentioned above with the following changes. Bacteria were lysed in Buffer-A (30 mM HEPEs pH 7.4, 50 mM KoAc, 2 mM MgOAc, 1 mM EGTA, 10% glycerol) supplemented with 2 mM PMSF, 5 mM DTT and protease inhibitor cocktail. Clarified lysate was loaded onto to Step-tactin column (GE Healthcare Life Sciences), washed with 20-30 column volumes wash buffer (Buffer A, 2mM PMSF, 5 mM DTT) and eluted with Buffer A supplemented with 5 mM DTT and 3 mM Desthiobiotin (SIGMA). Peak fractions were combined and further purified via size exclusion chromatography on a Superdex 200 (GE Healthcare Life Sciences) with degassed dialysis Buffer-A supplemented with 5mM DTT at 0.5 ml/min. Fractions corresponding to monomers/dimers of sfGFP-BICD2 (25-400 aa) were collected.

### Immunoprecipitations and binding assays

GST pull-downs were performed by incubating purified GST tagged proteins (5 µg) with 20 μl dry volume of glutathione agarose 4B beads (Thermo Scientific) for 1 h under rotation at 4 °C. Beads were pelleted by centrifugation at 500 xg for 1 minute and washed three times with GST pull down buffer (50 mM Hepes pH 7.4, 150 mM NaCl, 2 mM EGTA, 10% glycerol, 0.1% Triton-X100, 2 mM PMSF, 5 mM DTT) and incubated in with His-tagged bait protein (5 µg) in 200 µl GST pull-down buffer under end-over-end rotation for 2 h at 4°C. Beads were washed 5 times with 500 µl of GST pull-down buffer and eluted in 40 μl of SDS loading buffer by boiling for 5min. Samples (5-15 µl from the eluate) were loaded into 10% SDS-PAGE gels and stained with Coomassie Brilliant Blue (SIGMA) or transferred to a nitrocellulose membrane. In GST pull-downs containing nucleotides GST pull downs buffer was supplemented with 10 mM MgCl_2_.

Dynein-dynactin binding assays were performed by incubating 5 µg of His-tagged SEPT9 protein or truncations with 10 µl dry volume of Ni-NTA agarose beads (Macherey Nagel) for 1h under rotation at 4 °C. Beads were pelleted by centrifugation at 500 xg for 1 minute and washed three times with Buffer-A supplemented with 2mM PSMF and 5 mM DTT and subsequently incubated with 20 µl of native dynein-dynactin purified from HEK-293 cells (as described above) in a total volume of 200 μl Buffer-A supplemented with 2 mM PSMF, 5 mM DTT and 0.1% NP40. Beads were incubated under end-over-end rotation for 2 h at 4 °C. Beads were pelleted and washed five times with Buffer-A supplemented with 2 mM PSMF and 5 mM DTT and eluted in 20 µl of SDS loading buffer by boiling for 5 minutes. Samples were loaded into 6% or 10% SDS-PAGE gels and stained with Coomassie Brilliant Blue (SIGMA) (5 µl from the eluate) or transferred to a nitrocellulose membrane (15 µl from the eluate).

For co-immunoprecipitations experiments HEK-293 cells were plated on 10 cm dishes at ∼40% confluency the day before transfection in antibiotic free medium. Cells were transfected with 2 µg plasmid DNA using Metafetene-PRO (Biontex) according to the manufacturer’s instructions and 2 plates were transfected per condition. After 6 h, medium was exchanged for complete DMEM containing 10% FSB and PSK. Cells were grown for 24 h before lysate preparation. Cells were washed twice with ice cold PBS and collected by scraping in lysis buffer (Buffer-A supplemented with 2 mM PMSF, protease inhibitors cocktail, 5 mM DTT and 0.2% NP-40). To ensure lysis, cells were incubated for 30 minutes at 4 °C rotating end-over-end. Lysate was cleared by centrifugation at 16.000 xg for 20 minutes at 4 °C (Eppendorf). Supernatant was collected and protein content was quantified by Bradford Spectrophotometry (Biorad). Equal micrograms of protein extracts were mixed with 2 µg of Rabbit anti-mCherry antibody (Abcam) for 6 h at 4°C under end-over-end rotation. Protein AG beads slurry (25 µl; Santa Cruz) were added and the lysates were further incubated overnight at 4 °C under end-over-end rotation. Beads were washed five times with 500 µl of lysis buffer, boiled in 25 µl SDS loading buffer and run on a 10% SDS-PAGE gel. Gels were subsequently transferred to a nitrocellulose membrane and processed for western blotting.

### Cell culture, transfections and treatments

COS-7 (ATCC: CRL-1651), HEK-293T (ATCC: CRL-3216) and HeLa-CCL2 (ATCC: CCL-2) were maintained in a humidified incubator 37 °C with 5% CO_2_ in high glucose Dulbecco’s Modified Eagle Medium DMEM (DMEM, SIGMA) supplemented with 10% fetal bovine serum (R&D Systems) and 1% Penicillin/Streptomycin/Kanamycin (SIGMA and GIBCO). Primary rat embryonic (E18) hippocampal neurons were obtained from the Neuron Culture Service Center (University of Pennsylvania) and cultured in Neurobasal medium as previously described (Karasmanis et al 2018). BSC-1 cells stably expressing GFP-tubulin and mCherry-LAMP2 (gift from Melike Lakadamyali, University of Pennsylvania) were maintained as described previously (Mohan et al., 2019). For immunofluorescence or live imaging experiments cells were seeded at density of 70-100,000 cells on 22 mM glass coverlips or 35mm glass bottom dishes (MatTeck) coated with 30 µg/mL type I bovine collagen (Advanced Biomatrix).

For all immunofluorescence experiments COS-7 were transfected with 0.25µg DNA using Lipofectamine 2000 for 24 h unless otherwise stated. Medium was exchanged 6h post-transfection. MitoTagRFP-septin constructs were transfected using 1 µg of DNA and in co-transfection experiments, 0.75 µg of MitoTagRFP constructs was combined with 0.25 µg of GFP or GFP-p50 encoding plasmids. For the inducible peroxisome trafficking assay in Fig. 2 A, cells were transfected with 0.25 µg GFP-SEPT9-FRB plasmid and 0.25 µg PEX-RFP-FKBP plasmid. Cells were induced with 1 µM rapalog (AP21967, Takara) for 45 minutes at 37 °C. Cells in Fig. 1 L were transfected with two plasmids of equal concentration (0.25 µg each). For SEPT9 depletion experiments (Fig. 5 C and 5, Fig. S1 G), COS-7 and HeLa cells were seeded on a 10 cm dish at ∼40%-60% confluency one day before transfection with 8 µg plasmid DNA using 24 µl of Lipofectamine. Medium was exchanged 6 h post-transfection. After 48 h, cells were trypsinized and replated on 22 mm glass coverlips (80,000 cells per well). Cells were fixed and assayed 72 h post-transfection. Primary rat hippocampal neurons (DIV2-4) were transfected with 0.4-0.5 µg of ratSEPT9_i1-mCherry plasmid combined with 0.4 µg of plasmid encoding LAMP1-mGFP or GFP-Rab5A using Lipofectamine 3000 (Karasmanis et al., 2018). Neurons were assayed 48h post transfection. Nocodazole treatment was performed by incubating cells with 33 µM nocodazole (SIGMA) for 3 h and 0.1% v/v DMSO was used as vehicle control. Acute oxidative stress was induced by incubating COS-7 cells with 300 µM sodium-(meta)arsenite (NaAsO_2_; SIGMA) for 30 or 120 minutes (Willett et al., 2017) and HeLa cells with 400 µM NaAsO_2_ for 1 h. For glucose deprivation experiments cells were incubated in glucose-free DMEM (GIBCO) containing dialyzed FBS (GIBCO) for 1 h.

### Immunofluorescence

Cells in Fig1A, B and Fig. 5A were fixed with warm 2% PFA (Electron Microscopy Sciences) in PBS containing 0.4% w/v Sucrose for 12 minutes at room temperature and quenched with 0.25% Ammonium Chloride (SIGMA) for 10 minutes. Cells were simultaneously blocked and permeabilized in 0.2% porcine gelatin (SIGMA) and 0.1 w/v % Saponin (Millipore) for 30 minutes at room temperature. Primary antibodies were incubated overnight at 4 °C and secondary antibodies were incubated at room temperature for 1 h in 0.2% porcine gelatin BSA and 0.1% Saponin in PBS. Primary rat hippocampal neurons were fixed and stained as previously described (Karasmanis et al., 2018).

Cells in Fig. 1 H, I, L, Fig. S1 C, E, Fig. 4G and Fig. 5 C, H were fixed with warm 3% PFA (Electron Microscopy Sciences) in PBS containing 0.4% w/v Sucrose for 15 minutes at room temperature and quenched with 0.25% ammonium chloride (SIGMA) for 10 minutes. Cells were subsequently blocked and permeabilized with 3% BSA (SIGMA) and 0.2 w/v % Saponin (Millipore) for 30 minutes at room temperature. Primary antibodies were incubated for 1h at room temperature or overnight at 4°C in 3% BSA, 0.2% Saponin in PBS. Secondary antibodies were incubated at room temperature for 1h in 3% BSA, 0.2% Saponin in PBS.

Cells in Fig. 1D were fixed and stained as described previously (Mohan et al., 2019) with the following modifications. Briefly, cells were fixed with 4% PFA (Electron Microscopy Sciences) in PBS for 20 minutes and 0.25% ammonium chloride (SIGMA) was used to quench background fluorescence. Cells were blocked and simultaneously permeabilized with 3% BSA (SIGMA) and 0.2% Triton X-100 (SIGMA) dissolved in PBS. Cells were incubated with primary and secondary antibodies in blocking buffer and were rinsed between antibody incubation with 0.2% BSA and 0.05% Triton X-100.

Cells in Fig. 2, Fig. S2 C, E, and Fig. 4 C, E, were fixed with warm 2% PFA in PBS containing 0.4% w/v sucrose for 12 minutes at room temperature and quenched with 0.25% ammonium chloride (SIGMA) for 10min. Cells were permeabilized in 0.1% Triton for 10 minutes at room temperature and blocked with 0.2% porcine gelatin in PBS. Primary antibodies were incubated overnight at 4 °C and secondary antibodies were incubated for 1 h at room temperature in 0.2% porcine gelatin in PBS.

### Western blotting

Samples were loaded onto 10% or 6% SDS-PAGE gels and transferred to a 0.45 µm nitrocellulose membrane (Amersham GE Healthcare) overnight at 4°C in Tris-Glycine buffer under constant voltage (30V). Membranes were blocked with 5% non-fat dry milk and 1% BSA in PBS for 1 h at room temperature. Membranes were washed with PBS-T (PBS/0.1% Tween 20) and incubated with primary antibodies in PBS-T buffer containing 2% BSA and 0.025% sodium azide for 2 h at room temperature or overnight at 4°C. Subsequently, membranes were washed with PBS-T and incubated with anti-mouse or anti–rabbit secondary antibodies (LiCor) diluted in antibody dilution buffer for 1 h at room temperature before scanning with an Odyssey infrared imaging system (Odyssey; LICOR).

### Iodixanol density gradients

For membrane fractionation experiments two subconfluent 15 cm dishes were trypsinized and washed twice with PBS. Cell pellet was resuspended in homogenization buffer (20 mM HEPES pH 7.4, 1mM EDTA) supplemented with 2 mM PMSF (SIGMA) and protease inhibitor cocktail (SIGMA). Cells were incubated on ice for 10 minutes and were homogenized by passing 10-15 times through a 22G1/2 needle. Cell breakage was confirmed microscopically by Trypan Blue stain (SIGMA). To restore isosmotic conditions equal volume of homogenization buffer containing 0.5M Sucrose (SIGMA) was added to the cell homogenate. Nuclei were pelleted by centrifugation at 1000 xg for 5 min at 4 °C. The resulting post-nuclear supernatant (PNS) was diluted in OptiPrep gradient media to a final concentration of 15% OptiPrep (SIGMA) in a total volume of 1.5ml. The PNS was carefully layered on top of a discontinuous OptiPrep step gradient (17%, 20%, 23%, 27%, 30%) prepped in descending concentrations. The gradient was centrifuged at 145,000 xg in a SW60 Ti swinging bucket rotor (Beckman) for 2 h at 4 °C. Fractions (350 μl) were collected from top to bottom.

### Microscopy

Fixed samples were imaged on a Zeiss AxioObserver Z1 inverted microscope equipped with a Zeiss 20X/0.8 dry objective, a 40X/1.2 NA water objective, a 63x/1.4 NA oil objective, a Hamamatsu Orca-R2 CCD camera and the Slidebook 6.0 software. Fixed cells in Fig. 1A, B and live cell in Fig. S2 A were imaged on a laser scanning confocal microscope (LSM700; Carl Zeiss), equipped with an environmental chamber and operated by the Zen software (Carl Zeiss). Optical sections were acquired according to Niquist criteria with a 63X/1.4 NA Oil objective (Carl Zeiss). The PureDenoise plugin was used post-acquisition to remove Poisson shot noise from confocal images (Fig. 1 A, B) using an automated, global noise estimation in the FiJi software.

Super-resolution structured-illumination microscopy (SIM) was performed in Fig. 1D using DeltaVision OMX V4 (GE Healthcare) with an Olympus 60X/1.42-NA objective and immersion oil with a refractive index of 1.514. Images were acquired using a 0.125 µm z-step, sCMOS pco.edge camera (custom version) and reconstructed with softWoRx software (Applied Precision Ltd.). Total internal reflection fluorescence (TIRF) imaging (20-60 frames per min) of live neurons was performed at 37°C using the TIRF module on the DeltaVision OMX V4 inverted microscope equipped with an Olympus 60X/1.49 objective and a temperature-controlled stage-top incubator.

Time-lapse imaging of COS7 cells in Fig. 1 E, F, and Fig. S1 A was performed with an inverted Olympus IX83 spinning disk confocal system equipped with a Yokogawa CSU 10 spinning disk, a motorized stage, solid state 405, 488, 561 and 640 nm laser lines, a Hamamatsu Orca Flash 4 CMOS camera and an environmental control chamber (OKOLAB). Images were acquired using a 60X/1.49 NA oil objective (Olympus) and the VisiView software. COS-7 cells were co-transfected with plasmid expressing -SEPT9_i1-mCherry under the weak PGK promoter and plasmids encoding LAMP1-mGFP or GFP-EEA1. Cells were imaged 24 hours post transfection at 37 °C and 5% CO2 in Fluorobrite DMEM media supplemented with 10% FBS and 1% PSK. Low expressing cells were identified and imaged at 3 second intervals for a total duration of 5 minutes.

Live imaging of hippocampal neurons (DIV 4-6) was performed 48 h post-transfection with the internal reflection fluorescence (TIRF) microscopy module of the DeltaVision OMX V4 system using a 60X/1.49 NA objective and temperature (37 °C) controlled environmental chamber. Cell media was replaced with phenol red-free Neurobasal media supplemented with 2% B27 (Invitrogen) and 30 mM HEPEs and dishes were sealed using ParaFilm. Axons were identified as the longest process/neurite and imaged at 2-4 minute intervals. Movies were displayed at the rate of one frame per second.

For live cell imaging of Mito-Spy labeled cells (Fig. S2 A), cells were incubated with 50 nM MitoSpy™ NIR DilC1 dye for 30 min at 37 °C. Cells were subsequently washed twice with imaging medium (FluoroBrite DMEM, GIBCO) and imaged in the environmental chamber (37 °C, 5% CO_2_) with the Zeiss LSM700 confocal laser-scanning microscopy in an environmental chamber (37 °C, 5% CO_2_).

### Image analysis and quantifications

All image processing and quantitative analyses were performed with the open-source software Fiji. Ratios of perinuclear to peripheral fluorescence intensity were derived by dividing the mean perinuclear intensity by the mean peripheral intensity using the formula (I_Perinuclear-_ BG/Area_Perinuclear_)/[(I_Total_-BG)-(I_Perinuclear_-BG)]/(Area_Total_-Area_Perinuclear_) with I denoting mean fluorescence intensity and BG corresponding to mean background intensity. Perinuclear area was defined as the area contained within a 40 µm diameter circle positioned around the cell nucleus. Sum intensities for each area were measured and background (BG) was subtracted. Mean background intensity was measured by quantifying fluorescence intensity in a circular extracellular region and multiplying it by a factor to equal total cell or perinuclear surface area under quantification. Cell perimeter was traced manually using the freehand selection tool.

Cumulative lysosome intensity was measure and plotted as recently described. (Starling et al., 2016). Cell perimeter was defined by thresholding GFP fill expressed from shRNA plasmids and corrected manually where necessary using the freehand selection tool. Cell area was scaled in 10% decrements and sum intensity (integrated density) was measured for each decrease. Background was subtracted using the subtract background function in Fiji (rolling ball radius=50). Intensity of each fraction was normalized by diving by the total cell intensity. Cumulative LAMP-1 intensity was plotted against the respective cell fraction. Curves were fitted using non-linear regression, second order polynomial being the preferred fitting model for our data. Statistical significance between different fitting models and different experimental conditions was assessed using the extra-sum-of-squares F test.

The ratio of retrograde to anterograde motility in neurons was derived by dividing the number of retrograde motility events of mCherry or mCherry SEPT9 positive Lamp1 vesicles by the number of anterograde motility events per axon, per 2 minutes of time-lapse movie. Each axon was imaged once.

Percent intensity of perinuclear mitochondria was derived by dividing the sum perinuclear intensity by the sum cell intensity using the formula [(I_Perinuclear-_BG)/(I_Total_-BG)] x 100. Percent peripheral intensity was calculated by subtracting perinuclear intensity from 100. Perinuclear area was defined as the area contained within a 40 µm diameter circle positioned around the cell nucleus. Mean background intensity (BG) measured by drawing a circular extracellular area and multiplyin it by a factor to equal total cell or perinuclear area and considered background intensity (BG). Cell perimeter was traced manually using the freehand selection tool.

Percentage of EEA1- and LAMP1-positive organelles with SEPT9 (Fig. 1 C, Fig. S1 J) were quantified in COS7 cell and neuronal axon images after applying a median filter. Puncta with signal above background levels and an over 30% surface area overlap with membrane marker were scored as positive. Percentage of SEPT9 positive organelles was calculated by dividing the number of SEPT9 positive by the total number of organelles per cell and multiplied by 100. In Fig. 1 C, lysosomes in contact with septin filaments were not included in the quantification. In Fig. S1B, percentage of mCherry-SEPT9_i1 puncta that overlap with LAMP1-mGFP or GFP-EEA1 was calculated in Fiji by creating masks of mCherry-SEPT9_i1 puncta in the 594 nm channel and overlaying them with the LAMP1-mGFP or GFP-EEA1. Non-overlapping puncta were deleted using the DeleteSelected_ROI2 macro created by Kees Straatman (University of Leicester) and overlapping puncta were counted as percentage of total per cell. This quantification was performed in the first frame of the time-lapse series acquired by spinning disk confocal microscopy. Images in Fig1E-F, FigS1A and Video S1 were bleach corrected in Fiji (histogram matching).

Distribution of peroxisomes and lysosomes in (Fig. 2 A, Fig. 5 D and I) was assessed visually. Peroxisome (Fig. 2 A) and lysosome distribution (Fig. 5 D) was assessed visually and classified into three categories: clustered, partially clustered or dispersed. Lysosome distribution in Fig. 5 I was binned into two categories based on the abundance of lysosomes at perinuclear or peripheral sites. Cells with peripheral lysosomal accumulations were classified as peripheral and cells with perinuclear but no peripheral lysosome clusters were classified as perinuclear.

Quantification of the fluorescent intensity of DIC (Fig. 2 E, Fig. 4 F) or MitoSpy dye (Fig. S2 A), the mitochondria channel was processed in Fiji with a fast Fourier transform-based band pass filter to eliminate noise (Giedt et al., 2016)and mitochondria masks were subsequently created using an auto-threshold algorithm. Fluorescence DIC or MitoSpy intensity was measured in the mitochondria masks. Background subtraction was performed with the “rolling ball” background subtraction tool using a ∼50 pixel averaging subtraction ball.

In Fig. 5 B, SEPT9-LAMTOR4 overlap was quantified by creating masks that correspond to SEPT9 and LAMTOR4 signal intensities and an overlap mask was derived and corrected using the watershed separation algorithm. The surface area (µm^2^) of each overlap and LAMTOR4 mask and the percentage of overlap per cell was derived by dividing the total surface area of the overlap masks by the total surface area of LAMTOR4 and multiplying by 100. Masks were created from thresholded images after applying a Laplacian filter followed by Gaussian blur filtering (2 sigma radius). Background was subtracted using “rolling ball” background subtraction (50 pixel ball radius).

Quantification of SEPT9 depletion (Fig. S1, H and Fig. 5 G) was performed in images of SEPT9-stained cells transfected with scramble and SEPT9 shRNAs. Cell periphery was outlined manually using free-hand selection and the GFP signal from the pGFP-Super expressing plasmids as a guide. Background was subtracted using a round ROI at the extracellular area close to the cell of interest.

All western blots and Coomassie-stained gels were quantified in Image Studio Lite (LICOR-Odyssey).

### Statistical analysis

Statistics were performed in GraphPad Prism 5.0. Data sets were tested for normal distribution or variance using the D’Agostino and Pearson normality test. Data exhibiting normal distribution were tested using Student’s *t* test when comparing two data sets, while a one-way analysis of variance (ANOVA) with a Bonferroni post-hoc test was used when comparing multiple data sets. For non-normally distributed data a Mann-Whitney *U* test was used when comparing two data sets and a Kruskal-Wallis test with a Dunn’s Multiple Comparison Test was used when comparing multiple data sets. A two-way analysis of variance (ANOVA) with a Bonferroni post-hoc test was used when comparing grouped data sets. Qualitatively binned datasets were analyzed using a Chi-square test. All graphs were generated in GraphPad Prism 5.0.

### Online supplemental material

Fig. S1 shows the specificity of the localization and effects of SEPT9 on lysosomes. Fig. S2 shows the localization of mitochondria and endogenous septin paralogs in cells that express mitochondria-targeted septins. Video 1 shows co-trafficking of SEPT9_i1-mCherry with LAMP1-mGFP in COS-7 cells. Video 2 shows LAMP1-mGFP traffic in the axon of a rat embryonic hippocampal neuron (DIV4) that expresses mCherry. Video 3 shows LAMP1-mGFP traffic in the axon of a rat embryonic hippocampal neuron (DIV4) that expresses rat SEPT9_i1-mCherry.

## Abbreviations

aa: amino acids;
DHC: dynein heavy chain;
DIC: dynein intermediate chain,
DLIC: dynein light intermediate chain;
EEA1: early endosome antigen 1;
ESCRT: early sorting complex required for transport;
G-domain: GTPase domain;
GTP: guanosine-5′-triphosphate;
GDP: guanosine diphosphate;
GTPγS: guanosine 5′-O-[γ-thio]triphosphate;
JIP-1: JNK-interaction protein-1;
LAMP1: lysosomal associated membrane protein 1;
LAMTOR4: late endosomal/lysosomal adaptor, MAPK and mTOR activator 4;
MVB: multivesicular body;
NTE: N-terminal extension;
ROS: reactive oxygen species;
SEPT: septin;
TSG101: tumor susceptibility gene 101.

## Acknowledgments

We thank Drs. Erika Holzbaur (University of Pennsylvania), Adam Kwiatkowski (University of Pittsburgh), Melike Lakadamyali (University of Pennsylvania), Trina Schroer (The Johns Hopkins University) and Zu-Hang Sheng (National Institutes of Health) for donating constructs and helpful advice. We are grateful to Dr. Sam Reck-Peterson and John Gillies (University of California San Diego, Howard Hughes Medical Institute) for *in vitro* motility assays of dynein activation. We also acknowledge Megan Radler, Brenna Doyle and Drs. Konstantinos Nakos and Eva Karasmanis for help and feedback on the manuscript. All microscopy was performed in Drexel University’s Cell Imaging Center. This work was supported with NIH/NIGMS RO1 grant GM097664 and R35 grant GM136337 to E.T.S.

The authors declare no competing financial interests.

## Author contributions

Ilona Kesisova designed and performed the majority of the experiments, analyzed data and assembled figures. Ben Robinson performed experiments and analyzed data. Elias Spiliotis initiated and directed the study, and wrote the manuscript with feedback from I. Kesisova.

