## Supplementary Material for "A septin GTPase scaffold of dynein-dynactin motors triggers retrograde lysosome transport"

### Supplemental Figures (Kesisova et al)

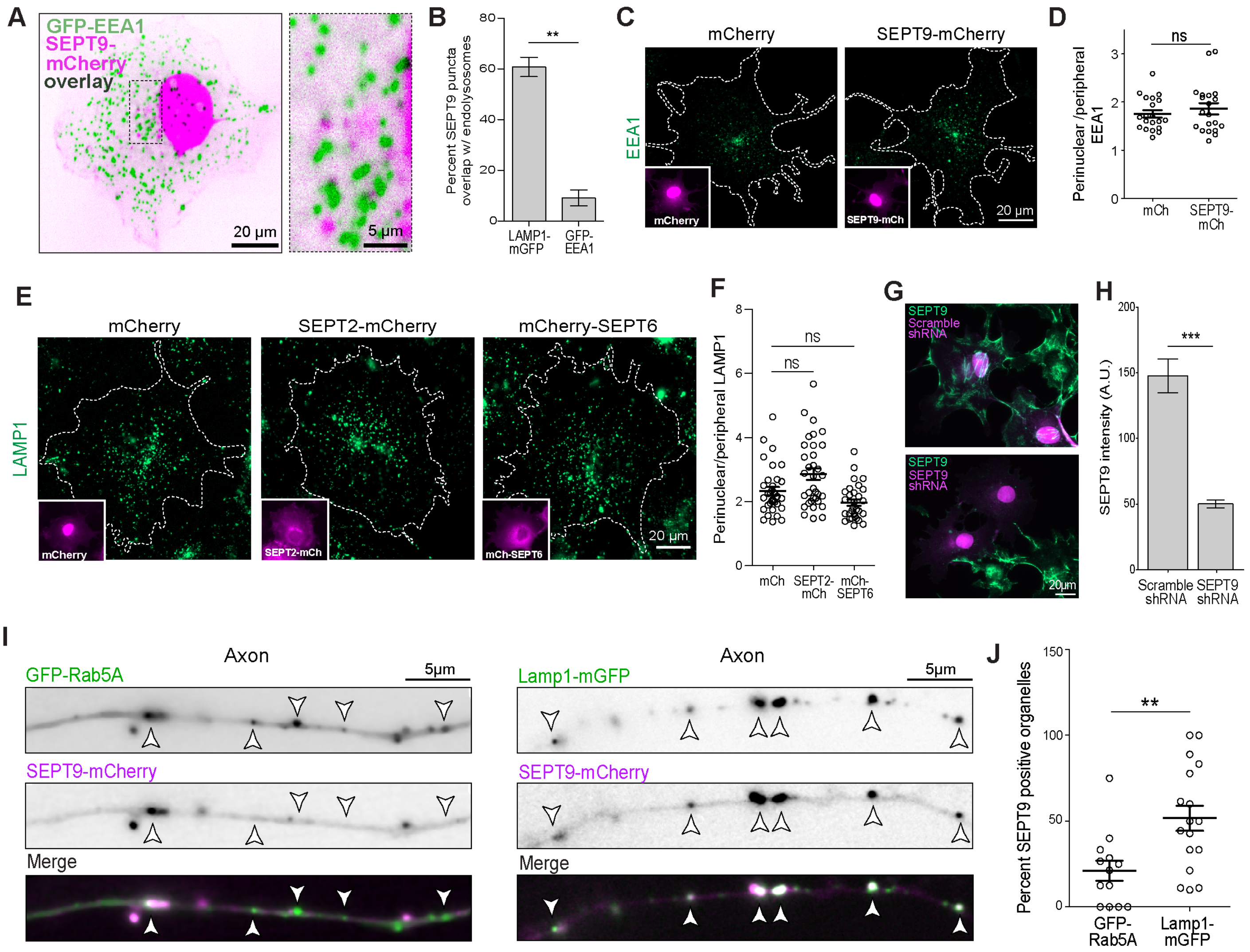


**Figure S1. Septin localization and effects on early endosomes and lysosomes.** (A) Spinning disk confocal image shows the distribution of SEPT9-mCherry, which was expressed under the PGK promoter, with respect to GFP-EEA1 in living COS-7 cells. Perinuclear area is shown in higher magnification. (B) Quantification shows the fraction (mean ± SEM) of SEPT9-mCherry puncta colocalizing with LAMP1-mGFP and GFP-EEA1 in COS-7 cells (*n* = 6). (C-D) Images (C) show the distribution of early endosomes (EEA1) in COS-7 cells that over-express mCherry or SEPT9-mCherry (insets). Quantification (D) shows the ratio (mean ± SEM) of perinuclear to peripheral fluorescence intensity of EEA1 per cell (*n* = 20). (E-F) Images (E) of LAMP1-stained COS-7 cells expressing mCherry, SEPT2-mCherry or mCherry-SEPT6 (insets). Plot (F) shows ratio (mean ± SEM) of perinuclear to peripheral fluorescence intensity of LAMP1 per cell (*n* = 30-34). (G-H) Images (G) of COS-7 cells transfected with plasmid DNA expressing GFP and scramble or SEPT9 shRNA after fixation and staining with anti-SEPT9. Bar graph (H) shows fluorescence intensity (mean ± SEM) of SEPT9 per shRNA-expressing cell (*n* = 27-35). (I-J) Images (I) of hippocampal neurons (DIV4) co-transfected with plasmid DNA expressing ratSEPT9-mCherry with GFP-Rab5A or LAMP1-mGFP. Plot (J) shows percentage (mean ± SEM) of GFP-Rab5A and LAMP1-mGFP organelles with SEPT9 per cell (*n* = 13 and 17 cells respectively). Ns, non-significant ($p > 0.05$); **, $p < 0.01$; *** $p < 0.001$.

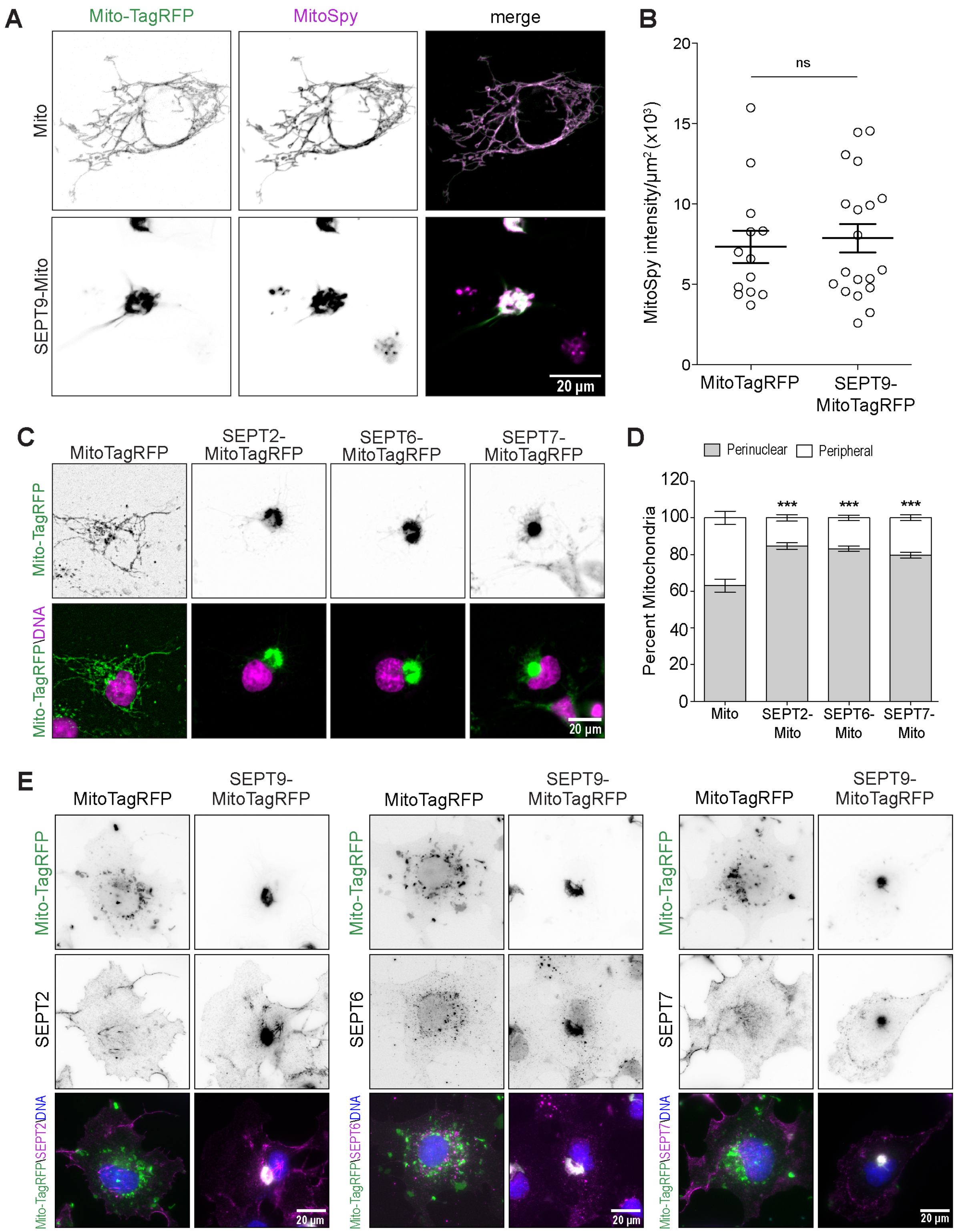


**Figure S2. Mitochondria-targeted septins induced.** (A-B) Images (A) show localization of MitoTagRFP- and SEPT9-MitoTagRFP -labeled mitochondria in living COS-7 cells, which were labeled with MitoSpy as an indicator of cell and mitochondria health. Plot (B) shows the fluorescence intensity (mean ± SEM) of MitoSpy per mitochondria (MitoTagRFP) surface area per cell (*n* = 13-19). (C-D) Images (C) show COS-7 cells transfected with MitoTagRFP (control) and SEPT2-, SEPT6- and SEPT7-MitoTagRFP after staining with DAPI. Bar graph (D) shows percentage (mean ± SEM; *n* = 20-28 cells) of total mitochondria fluorescence intensity in the perinuclear and peripheral cytoplasm. (E) Images show COS-7 cells transfected with MitoTagRFP or SEPT9-MitoTagRFP and stained with antibodies against SEPT2, SEPT6 and SEPT7. Ns, non-significant ($p > 0.05$); *** $p < 0.001$.

**Video 1**: COS-7 cells were co-transfected with LAMP1-mGFP and under the control of a weak PGK promoter, and imaged every 3 sec by spinning disk microscopy. Arrows point to SEPT9 positive lysosomes moving retrogradely. SEPT9_i1-mCherry is pseudo-colored in inverted magenta and LAMP1-mGFP is pseudo-colored in inverted green. Video display rate is at 5 frames/sec. Related to Fig. 1.

**Video 2**: Rat embryonic hippocampal neurons (DIV-4) were co-transfected with mCherry and LAMP1-mGFP, and imaged with shallow-angle TIRF microscopy every second for two minutes. mCherry is pseudo-colored in magenta and Lamp1-mGFP is pseudo-colored in green. Video display rate is at 15 frames per second. Neuronal cell body (soma) is located to the left of the axonal segment of the video (i.e., particles move retrogradely to the left and anterogradely to the right). Scale bar, 5 μm. Related to Fig. 1.

**Video 3**: Rat embryonic hippocampal neurons (DIV-4) were co-transfected with rat SEPT9_i1-mCherry and LAMP1-mGFP, and imaged with shallow-angle TIRF microscopy every second for two minutes. Rat SEPT9_i1-mCherry is pseudo-colored in magenta and LAMP1-mGFP is pseudo-colored in green. Video display rate is at 15 frames per second. Neuronal cell body (soma) is located to the left to the axonal segment of the video (i.e., particles move retrogradely to the left and anterogradely to the right). Scale bar, 5 μm. Related to Fig. 1.
